# A cognitive map for value-guided choice in ventromedial prefrontal cortex

**DOI:** 10.1101/2023.12.15.571895

**Authors:** Sebastijan Veselic, Timothy H. Muller, Elena Gutierrez, Timothy E. J. Behrens, Laurence T. Hunt, James L. Butler, Steven W. Kennerley

## Abstract

The prefrontal cortex is crucial for economic decision-making and representing the value of options. However, how such representations facilitate flexible decisions remains unknown. We reframe economic decision-making in prefrontal cortex in line with representations of structure within the medial temporal lobe because such cognitive map representations are known to facilitate flexible behaviour. Specifically, we framed choice between different options as a navigation process in value space. Here we show that choices in a 2D value space defined by reward magnitude and probability were represented with a grid-like code, analogous to that found in spatial navigation. The grid-like code was present in ventromedial prefrontal cortex (vmPFC) local field potential theta frequency and the result replicated in an independent dataset. Neurons in vmPFC similarly contained a grid-like code, in addition to encoding the linear value of the chosen option. Importantly, both signals were modulated by theta frequency – occurring at theta troughs but on separate theta cycles. Furthermore, we found sharp-wave ripples – a key neural signature of planning and flexible behaviour – in vmPFC, which were modulated by accuracy and reward. These results demonstrate that multiple cognitive map-like computations are deployed in vmPFC during economic decision-making, suggesting a new framework for the implementation of choice in prefrontal cortex.

## Main text

The prefrontal cortex (PFC) is fundamental for learning and choice^1–10^. A dominant idea in economic decision-making has been that value is represented in a common currency format, which facilitates efficient action selection^1,5,11^. While these studies have led to great progress towards understanding PFC’s neural code in relatively simple and overlearned decision contexts, humans and animals often face novel choices where they must infer or construct value based on previous experience^12,13^. Here, we investigated whether making choices of this kind requires a new framework for representing choice options within a value space; one in which the representation of choice task structure becomes crucial.

In contrast to research on the PFC, research on the medial temporal lobe (MTL) has investigated representations of structure and representations supporting inference^14^. In the MTL, cognitive maps encoding the relationships between entities in the world are built, supporting flexible behaviour^15,16^. Two key neural substrates allow for this: the ability to infer vectors between different locations using a grid-like code^14^, and planning-related signals observed during replay and coinciding with sharp wave ripples (SWR)^17,18^. Grid cells represent the structure of space^19–21^ and such structural representations are thought necessary for inferential choices that go beyond direct experience^14,16,21^. Relatedly, ripples in MTL may reflect planning signals during model-based reinforcement learning^18^, as place cells encode sequences of locations during replay^17^, which may underlie our ability to compositionally bind information and help facilitate novel choice^22^.

While investigating cognitive map-like representations and computations has historically focussed on physical space and the MTL, such signals have now been found in abstract spaces and outside MTL^12,13,15,23–27^. fMRI work in (non) human primates has implicated the ventral and medial parts of PFC (mPFC, vmPFC)^12,13,15^, perhaps due to their strong anatomical links with the MTL^28,29^. This suggests the computations supporting inference, novel choice, and planning in spatial navigation may be a general neural mechanism implemented in the brain; however, the link to choice has not yet been shown. Motivated by recent findings in non-human primate fMRI showing a grid-like code in a value space as subjects passively navigated between trials^12^, we asked whether the same neural code is present during choice itself, which would suggest it is used for navigation between possible (choice) locations in abstract space.

Here, we demonstrate the presence of a grid-like code at choice, defined by the trajectory between choice options in a two-dimensional (2D) value space. This grid-like code is present in local field potential (LFP) and single neurons in vmPFC, suggesting such a code is used for making choices in a similar fashion to navigating routes between locations in physical space. In addition, we show that while this code is stable over the same stimulus sets, it realigns over new stimulus sets, analogous to grid realignment observed in physical space, deepening the parallel between spatial and abstract map-like representations. Finally, we report the first evidence of ripples in non-human primate vmPFC and show these ripples are present primarily at choice and outcome, suggesting their role in binding choices and outcomes. Jointly, these results suggest the neural code in vmPFC underlying economic decision-making bears resemblance to the well-characterised representations of cognitive maps in the MTL that support inferential choices in space. This bridges two seemingly disparate fields – one traditionally focused on space and memory in the MTL, and one focussing on value and PFC – in a more unified perspective.

### Behaviour and chosen value representations in PFC

Two male rhesus macaques (Macaca mullata) made decisions between two options (Figure 1A), where an ‘option’ had to be constructed compositionally from two previously learned cues (images) that never formed an option during training (See Methods). A set of five cues mapped to one out of five reward probability levels, while another set of five cues mapped to one out of five reward magnitude levels. Subjects’ choice accuracy was above chance (Figure 1BC). We first looked for canonical value signals in single neurons across the four regions we recorded from (see Supplementary Figure S1): anterior cingulate cortex (ACC, n = 198), dorsolateral prefrontal cortex (dlFPC, n = 156), orbitofrontal cortex (OFC, n = 195), and vmPFC (n = 160), in line with how data from such experiments are typically analysed. Neurons in ACC, dlPFC, and OFC significantly encoded chosen value (Figure 1DE) and chosen value difference (Figure S2) in a 300-millisecond window before subjects initiated their choice. The signal was overall strongest in ACC neurons compared to the other three regions, thereby matching similar studies^3,30^. In contrast, we found no evidence for a chosen value or chosen value difference signal in vmPFC neurons during the same time period. This highlighted a frequent divergence between fMRI research, where choice-related value signals in vmPFC are commonly found^12,31–33^, and findings from electrophysiology, where choice-related activity is often weaker or absent relative to neurons in other regions^34–36^.

**Figure 1:**
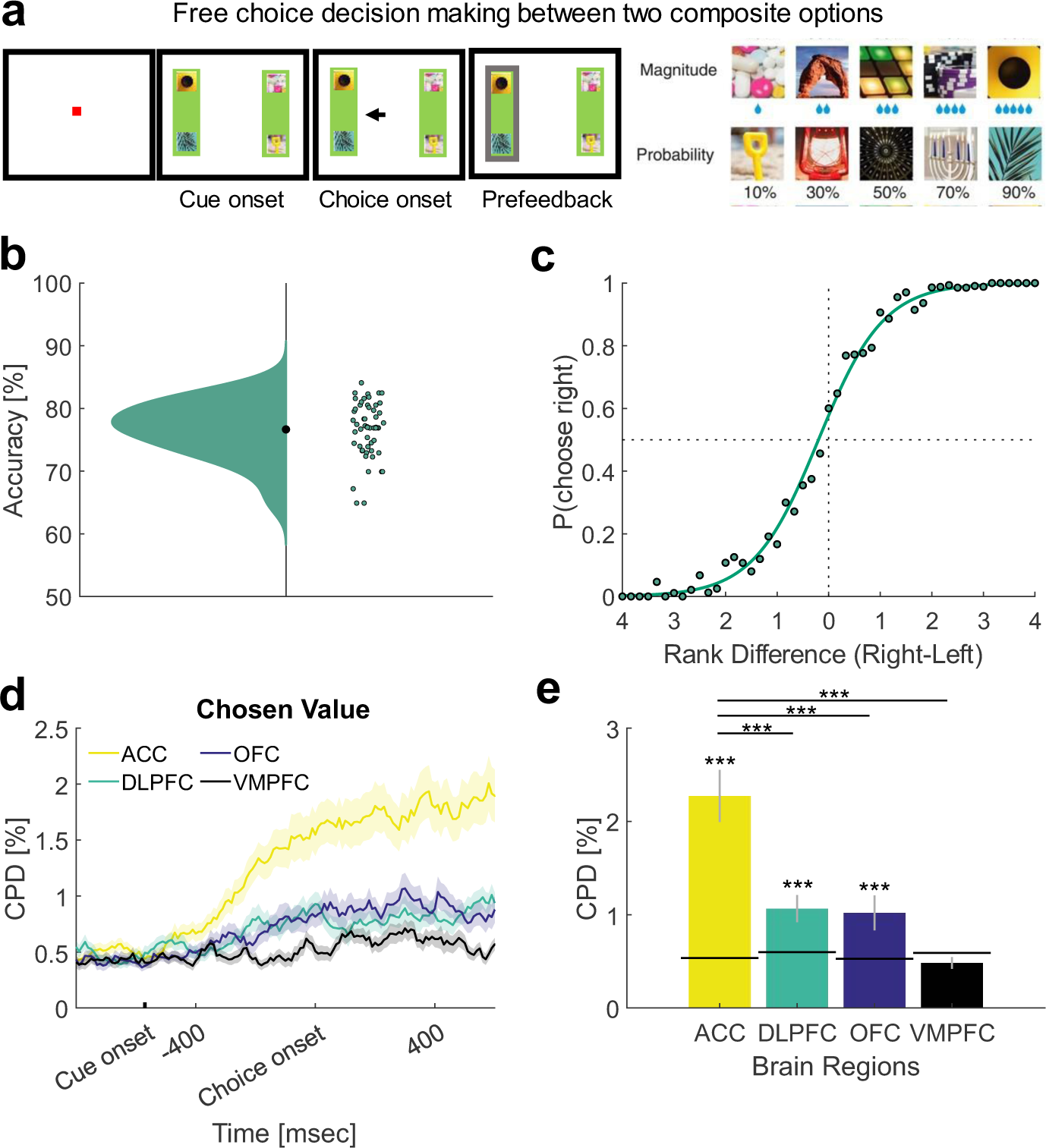
Task design and canonical value representations. **a)** Subjects were simultaneously presented with a left and right choice option, each comprising two cues (images). Once the cues were presented (Cue onset), subjects were free to saccade to the cues and choose their preferred option by moving the joystick towards it (Choice onset). The choice was confirmed by the highlighting of the selected option (Prefeedback), followed by reward delivery. The set of ten unique cues, each representing one of five reward magnitude or reward probability levels, was swapped every three to four recording sessions. Each stimulus set had ten new unique images, which represented the same set of five reward magnitude and reward probability levels (right panel). **b)** Accuracy (i.e., choosing the option with the highest expected value) across sessions**. c)** Psychometric curve across all trials**. d)** Chosen value coefficient of partial determination (CPD) for individual brain regions across the choice epoch. The thick line represents the mean response across the population within the brain region, the shaded area represents the standard error of the mean (SEM) across neurons within the brain region. Cue onset was approximately 560 msec before choice onset (mean reaction time across sessions). **e)** Mean CPD for the same regressor as in d) for individual brain regions within a 300-millisecond time window before subjects initiated their choice. The black line denotes the 95^th^ percentile of a null distribution for visual purposes, obtained through permutation testing (1000 permutations). *** p < .001. Error bars represent SEM across neurons within the brain region.

### A cognitive map of value space in vmPFC emerges at choice

While we did not find canonical value signals in vmPFC, we hypothesised the presence of map-like representations, based on previous work showing a grid-like code within vmPFC and adjacent (medial PFC) regions^12,13,15^. We predicted the cues representing individual attributes would be composed into options and mapped onto a reward magnitude by reward probability value space (Figure 2A, left panel), where a choice option corresponds to a location in 2D space. Each choice between two options reflects a trajectory between two locations in this 2D space. This is analogous to vector-based navigation in physical space^14^. We tested the hypothesis that subjects construct a cognitive map of value space using a grid-like code. We did this by looking for periodic modulation of neural activity predicted to arise from firing rate properties of grid cells^37,38^. Specifically, we looked for a grid-like code in local field potential activity in the theta frequency, based on previous work in physical space^39^. Such hexadirectional analyses rely on a pattern of signal where trajectories that are aligned with the grid field elicit stronger neural activity compared to those that are misaligned (Figure 2A, middle and right panel).

**Figure 2:**
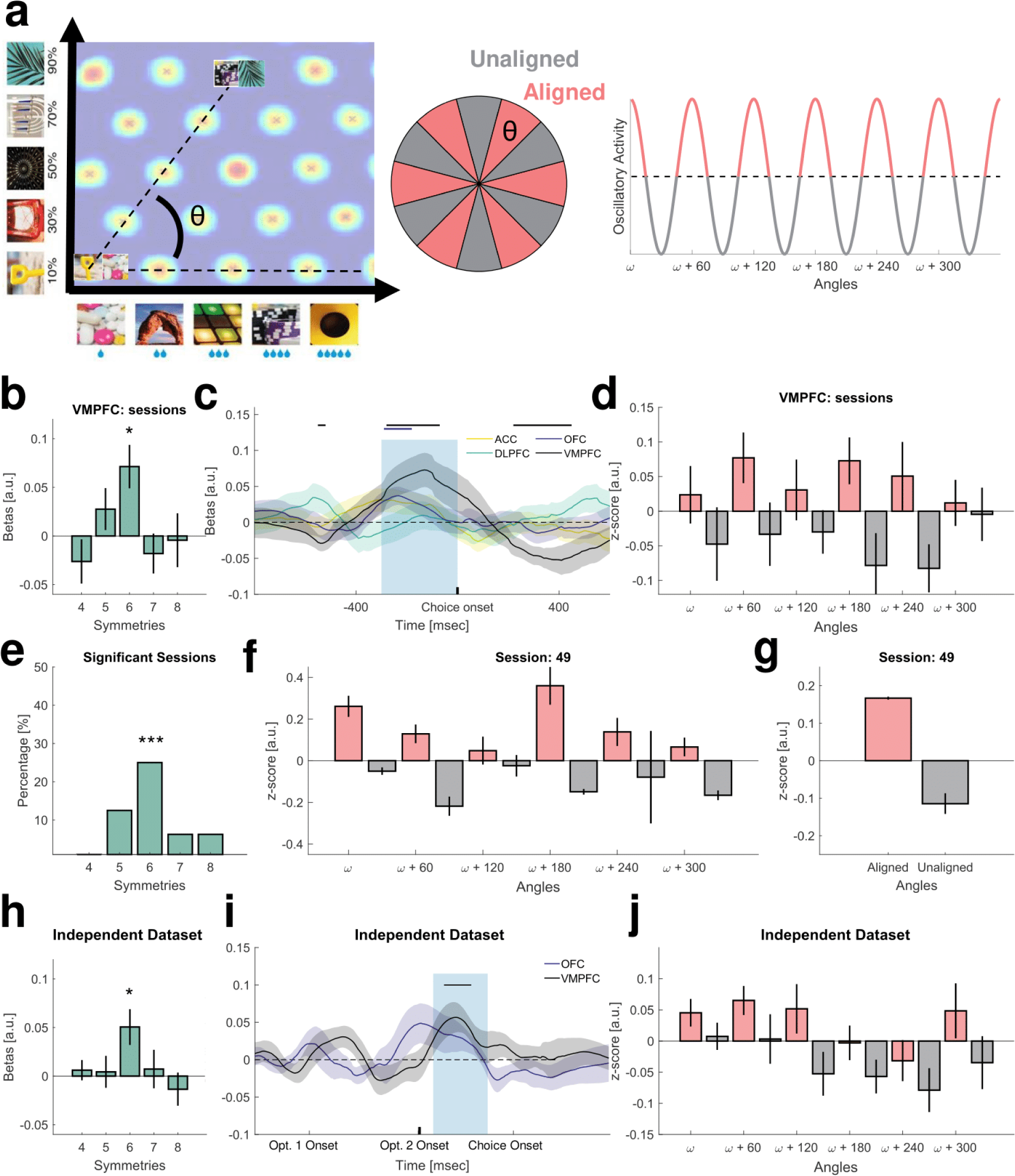
The value space is represented with a grid-like code at choice in vmPFC. **a)** Left panel: A value space is organised along the reward magnitude and reward probability values used for choice. Within this value space, the left and right options are embedded as “locations”. A trajectory or navigation angle can be computed between each pair of possible locations. Middle & right panel: Navigation angles falling along the grid field of a hypothetical grid cell (aligned) are predicted to elicit stronger oscillatory activity compared to angles that do not fall along the grid field (unaligned). **b)** Significant hexadirectional (sixfold) modulation in vmFPC but not control symmetries across recording sessions. Each session represents the average of several channels recorded within that session. See also Figure S3A. Error bars represent SEM across sessions (n = 16). * pBonferroni < .05, corrected for symmetries. **c)** Time course of hexadirectional (sixfold) modulation in vmPFC and other brain regions. Blue shading denotes the original time window in Figure 2B and Figure 1E. Lines above brain regions denote significant hexadirectional (sixfold) encoding at p < .05. See also Figure S3B. Error bars represent SEM across sessions within brain region (dlPFC = 32, OFC = 34, ACC = 41). **d)** Sixfold periodicity in vmPFC across sessions as predicted by Figure 2A, right panel. Error bars represent SEM across sessions. **e)** Percentage of significant sessions for individual symmetries obtained through permutation testing (n = 1000) by comparing session signal averages to the 99^th^ percentile of a null distribution. **f)** Sixfold periodicity for an example session. Error bars represent SEM across channels within that session (n = 3). **g)** Average signal for aligned compared to unaligned trials. Error bars represent SEM averaged across recorded channels (n = 3) within that session. **h)** Significant hexadirectional (sixfold) modulation in vmPFC but not control symmetries in an independent dataset occurring 100-500 msec after Option 2 onset. See Figure S5 for the task description. * p < .05. Error bars represent SEM across sessions (n = 11). **i)** Time course of hexadirectional (sixfold) modulation in vmPFC and another brain region (OFC) in an independent dataset. Error bars represent SEM across sessions within brain region (OFC, n = 8). **j)** Sixfold periodicity in vmPFC across sessions in an independent dataset. Error bars represent SEM across sessions.

We first regressed out all canonical value signals and reaction times, ensuring any obtained signals would not be confounded by them (see Methods). In the resulting residuals we found a cognitive map of value space, represented with a grid-like code in vmPFC theta frequency in a 300-millisecond window before subjects initiated their choice (Figure 2B, t(15) = 3.20, p = 0.006^1^) across recording sessions. The effect appeared before subjects made their choice (Figure 2C) and had distinct sixfold periodicity (Figure 2D), as in previous work investigating grid-like coding^15,38,39^. In contrast, there was no grid-like code in control LFP frequencies (Figure S3C), as in ref^39^. Individually, 25% of sessions (4/16) showed significant 6-fold modulation (Figure 2E, Binomial test, p < .001, see Figure 2FG for an example individual session). We found no grid-like code in ACC (t(40) = 1.54, p = 0.13) or OFC (t(33) = 1.73, p = 0.09), despite previous reports^15,40^, nor any grid-like signal in dlPFC (*t*(31) = 0.12, p = 0.182; see Figure S3ABD for regional comparisons).

In additional analyses, we observed two suggestive trends. The first trend implied the grid-like code in vmPFC became stronger as a stimulus set became more familiar (Figure S3EFG), perhaps suggesting the cognitive map is constructed during choice and further consolidated during sleep by hippocampal sharp wave ripples^41^. The second trend suggested the neural geometry of the cognitive map in vmPFC is better explained by a veridical compared to a distorted representation of the cognitive map (Figure S4).

Finally, to demonstrate the robustness of our result, we replicated the main effect from Figure 2B using the same analysis approach in an independent dataset that is currently being collected (Figure 2H, t(10) = 2.74 p = 0.02), see Figure S5 for a task explanation of the independent dataset). The effect similarly occurred before choice (Figure 2I) and exhibited sixfold periodicity (Figure 2J).

These results show subjects construct a cognitive map of value space with a grid-like code in vmPFC, echoing previous reports^12,13^, and suggest how it may be used for choice. Subjects construct a cognitive map by compositionally binding cues into options and embed these options as “locations” in a value space spanning choice-relevant attributes. Because this embedding occurs at choice on a trial-by-trial basis, this allows for computing navigation trajectories between choice locations in the value space. Crucially, the navigation trajectory between choice options and the length of that trajectory within a value space allows for computing the decision variable and thus allows for optimal choice (Figure 2A). This suggests a grid-like code may be used for the choice process itself, where it could facilitate inference of novel choices^13,26^ analogous to how grid cells facilitate inference of spatial shortcuts^14^.

### Grid orientations realign across stimulus sets but are stable within stimulus sets

To deepen the parallels with spatial cognition, we next tested whether the observed grid-like code was consistent across sessions and stimulus sets^15^. Grid cell grid orientations are typically stable within the environments they are recorded in but realign their grid fields across different environments^42^. Therefore, if the signal we observed indeed arose from populations of grid cells, this would predict a systematic difference in grid orientations across sessions, but only when the stimulus sets (i.e. “environment”) differed. Our study was unique in that every few sessions the stimuli denoting reward magnitude and reward probability values were changed (Figure 3A). Subjects had to learn these in a separate experiment interleaved with the choice task reported here. This meant identical value spaces (i.e. identical task structure) were decorrelated from sensory properties of stimuli subjects observed, analogous to having different sensory experiences in different spatial environments with identical underlying spatial structure^42^. As predicted, when we used grid orientations from one session to estimate a grid-like code in different sessions, we only observed a consistent grid-like code within (Figure 3B, t(8) = 4.77, p = .0014^2^) but not across stimuli sets (Figure 3C, t(15) = 0.06, p = .95).

**Figure 3:**
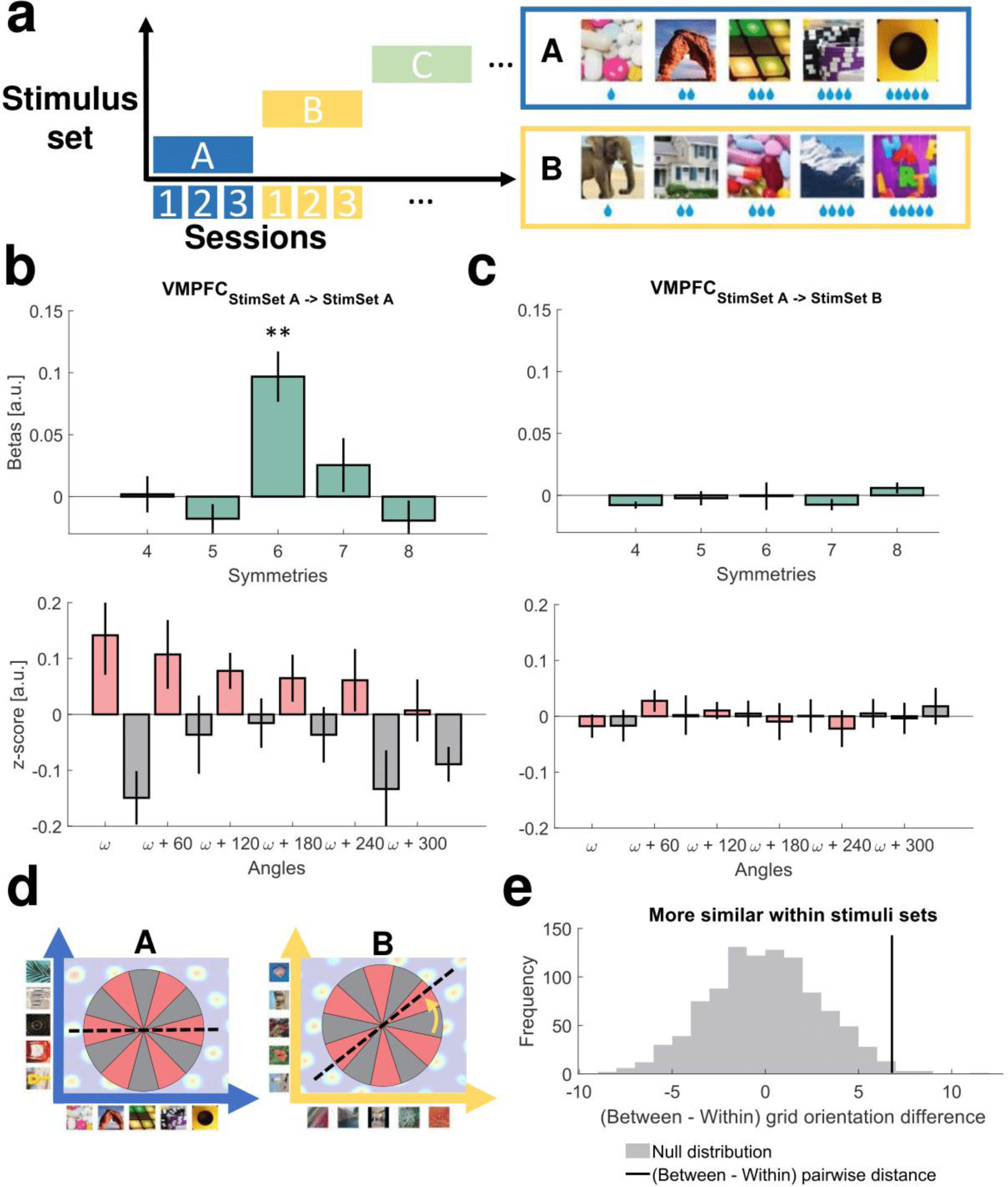
Grid orientations are stable within stimulus sets but realign across stimulus sets. **a)** The stimulus set denoting reward magnitude and reward probability was swapped every few sessions. The stimuli denoting the five reward magnitude and reward probability levels were not repeated and did not correlate with one another across sessions. **b)** grid orientations are stable across session comparisons within stimulus sets as observed by hexadirectional (sixfold) modulation in vmPFC. ** pBonferroni < .01. Error bars represent SEM across session comparisons within stimulus sets (n = 9). **c)** grid orientation is not stable across session comparisons across stimulus sets. Error bars represent SEM across session comparisons across stimulus sets (n = 16). **d)** schematic representation of the observed effect – the orientation of a putative grid field with respect to the underlying value space rotates from one environment to another environment. **e)** The average difference in grid orientations from different sessions with shared stimulus sets was smaller than the average difference in grid orientation from different sessions with different stimulus sets. The black line denotes the empirical value obtained for the difference of between-within grid orientation distances. The gray histogram denotes the shuffled null distribution (n = 1000 permutations).

We further demonstrated differences in grid orientations were not due to noise by computing grid orientation angle distances between sessions, comparing within vs. between stimuli set sessions. Average distances between grid orientations within stimuli sets were significantly smaller compared to ones across stimuli sets (Figure 3DE, z = 2.42, p < .01). The result in neither Figure 3B nor Figure 3E was not explained by potential within-session grid orientation correlations across channels as these were removed from the analysis beforehand. It was also not explained by time differences of pairwise comparisons (i.e. degree of exposure to a stimulus set) on which the distances were computed as a control analysis (Figure S6).

Overall, these results are the first demonstration of non-spatial, or abstract, grid realignment. Given grid cell realignment occurs when generalising across spatial contexts^42^, our results suggest similar mechanisms may be at play in non-spatial generalisation. This deepens the connection between known computational properties of grid cells measured in rodents during spatial navigation^42^ and our understanding of grid-like codes in non-spatial environments of other species. It also further provides evidence that the hexadirectional analysis used to estimate grid-like codes likely measures underlying grid cell activity, and hence supports the idea of prefrontal cortex containing grid cells constructing cognitive maps of abstract spaces.

### A grid-like code in vmPFC neurons and its theta phase-dependency

Having established a cognitive map of value space exists in the vmPFC LFP signal, we next sought evidence for such a representation in vmPFC neurons. Despite multiple studies demonstrating population-level grid-like codes in BOLD and LFP activity^12,13,15,38,39^, it has not been shown whether neurons represent information with a grid-like code as well.

Similar to cells in the hippocampal formation and OFC^43,44^, we observed neurons in vmPFC exhibit theta modulation (Figure 4ABC, F(9, 159) = 8.87, p = 4.20-13, see also Figure S7ABC) – firing most at theta troughs and firing least at theta peaks. We therefore isolated our hexadirectional analyses in vmPFC neurons to specific phases of several theta cycles using grid orientations from vmPFC channels (Figure 4D). This approach yielded a significant grid-like code across vmPFC neurons, but only at the trough of the average theta phase (Figure 4E, t(159) = 3.08, p = 0.002^3^), which was when cells fired the most (Figure 4A, right panel).

**Figure 4:**
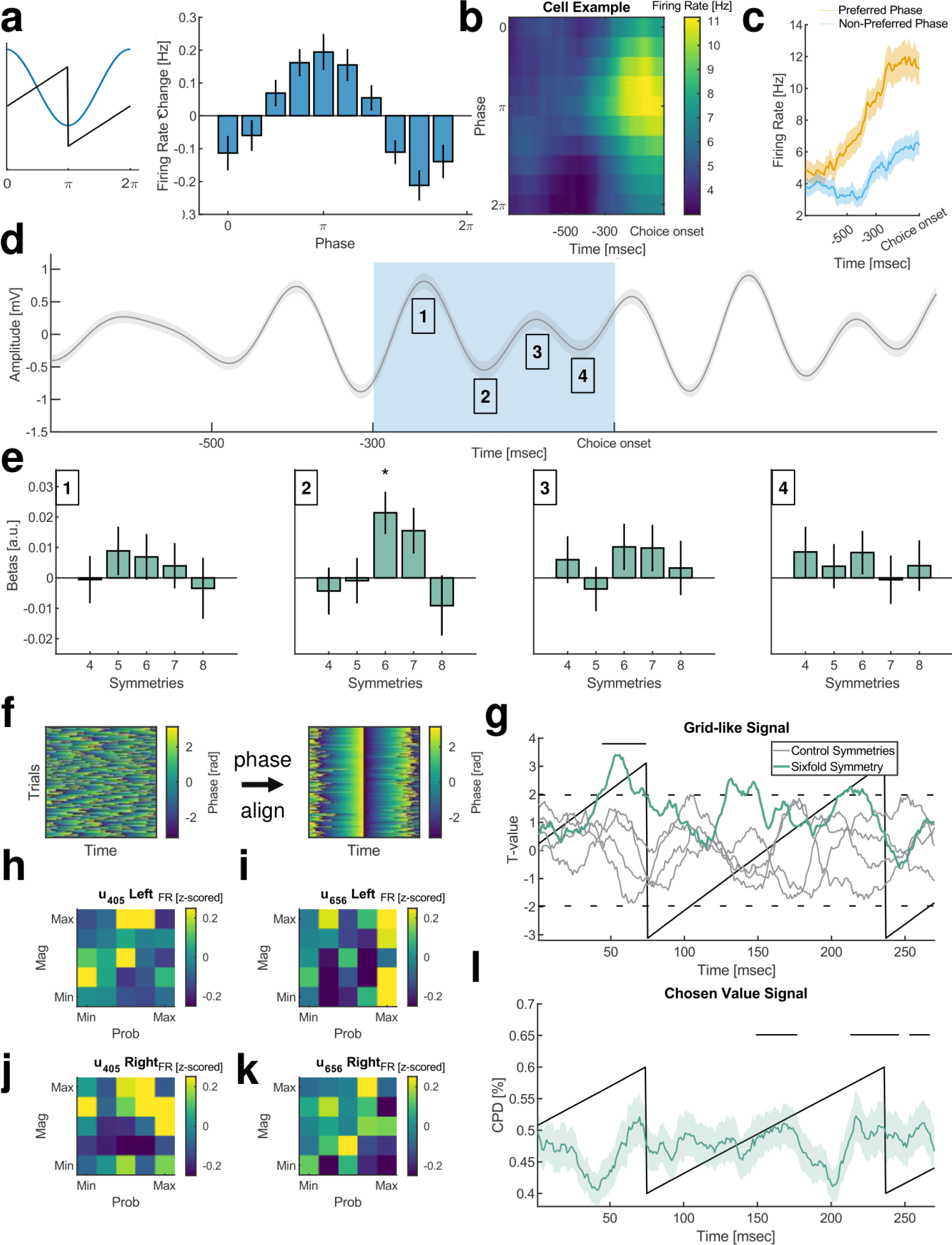
vmPFC neurons are theta modulated and maintain a grid-like code and chosen value code in separate theta cycles. **a)** Left panel: a sample theta cycle (blue) with its corresponding phase (black) showing the cycle–phase convention used throughout this figure. Right panel: vmPFC neurons were modulated by theta phase, firing most at theta troughs and firing least at theta peaks. Error bars represent SEM across vmPFC neurons. **b)** An example neuron plotted across all theta phases; representing how neurons responded on average (see S5BC for different patterns). **c)** The same example neuron from b) plotted at the Preferred and Non-Preferred phase – determined by t-tests across all cells (see Figure S7A). The neuron increased its firing closer to choice at a higher rate when indexed at theta troughs (Preferred) compared to theta peaks (Non-Preferred). Error bars represent SEM across trials within the session (n = 228). **d)** Average theta LFP across all vmPFC channels. The blue shading denotes the original time window in Figure 2B and Figure 1E. The numbers denote seed windows centred on theta throughs and peaks occurring within this time window. Error bars represent SEM across all vmPFC channels (n = 97). **e)** Significant hexadirectional (sixfold) modulation in vmPFC neurons at theta trough. * pBonferroni < .05. Error bars represent SEM across vmPFC neurons. **f)** Theta phase distribution across trials and time for an example channel before (left) and after (right) aligning it in time to one theta cycle. **g)** A grid-like code (green line), like in e), can be observed within one theta cycle before choice. The strength of the grid-like code is expressed as a t-value obtained from a t-test against zero across all cells using estimates from the hexadirectional analysis. Control symmetries are represented with the gray line. The dotted black lines represent t-test significance thresholds (significant t-test against zero) for visual purposes. The black bar denotes significance in cluster-based permutation testing (exceeding the 97.5^th^ percentile of the length-corrected null). **h)** Firing activity of an example neuron which exhibited a high grid-like code, plotted in the reward magnitude by reward probability space used to test for a grid-like code and averaged for the left option. The firing activity was averaged over the time period from Figure 4G where a significant grid-like code was found. **i)** firing activity of another example neuron with a high grid-like code, averaging over the left option. **j)** the same neuron from h) averaged for presentations of the right option. **k)** the same neuron from i) averaged for presentations of the right option. **l)** a chosen value signal can be observed at a different, subsequent, theta cycle before choice. The black bars denote significance in cluster-based permutation testing (exceeding the 97.5^th^ percentile of the length-corrected null). Error bar represents SEM across vmPFC neurons.

To further validate this signal was specific to theta troughs, we aligned theta phases in time on a trial-by-trial basis for each channel (see example for one channel in Figure 4F) and shifted the firing rates of vmPFC neurons based on their corresponding channels. This ensured that across neurons, neuronal firing at different timepoints was precisely synchronized to the theta phase of the channel on which a neuron was recorded. This analysis similarly revealed a grid-like code in vmPFC neurons at one theta trough (Figure 4G), which was temporally localized (Figure S7G) to the approximate time period as the theta trough labelled **‘2’** in Figure 4DE. This temporal pattern suggests the observed grid-like code occurred predominately within one theta cycle before choice – however – it remained weakly significant when we averaged over all theta troughs occurring in the 300-millisecond window before subjects initiated their choice from Figure 2B (Figure S7D, t(159) = 2.26, p = 0.025). When we plotted vmPFC neurons with high grid-like coding estimates (Figure 4HIJK), their rate maps had irregular firing fields, exhibiting a higher firing rate for multiple non-neighbouring states in the value space.

Because we found a theta phase-dependent grid-like code in vmPFC neurons, we tested whether this may explain the lack of chosen value signals in vmPFC in Figure 1DE, given previous work^45^, and considering other recorded regions encoded this information. Using the theta phase-aligned neuronal firing rates mentioned above, we indeed observed the chosen value signal occurring in a theta phase-dependent manner (Figure 4L). vmPFC neurons encoded chosen value at the trough of the theta cycle following the trough of the theta cycle containing a grid-like code; suggesting the cognitive map must first be constructed, and the relevant navigation angle computed, before estimates of chosen value can be represented. As an additional test, we used the same seed windows from Figure 4D and replicated this observation whereby vmPFC neurons weakly encoded chosen value at the subsequent theta trough (Figure S7E; corresponding to the time period labelled as ‘**4**’ in Figure 4DE) but not the one before. Across neurons, the chosen value signal did not correlate with the strength of the grid-like code (Figure S7F, S7I), suggesting neurons supporting cognitive map-like representations are independent from neurons representing chosen value. This implies both neuronal subpopulations may interact with one another to facilitate optimal choice.

More broadly, these results show vmPFC neurons may employ a theta phase-dependent neural code, which could be a possible explanation for the divergence between vmPFC findings in fMRI^12,31–33^ and electrophysiology^34–36^. Furthermore, finding a cognitive map of value space in vmPFC neuronal activity strongly suggests this region could contain grid cells.

### Sharp wave ripple events are present in vmPFC

Considering subjects had constructed a cognitive map for value that was used during choice, we looked for evidence of other MTL cognitive map-related computations in vmPFC. Another such phenomenon are sharp wave ripples (SWR) – and associated replay – in awake animals occurring at choice points^17,41,46^. SWR-associated replay is suggested to be a neural mechanism for the retrieval of information^41,47,48^, and model-based planning^18^.

We used a ripple detector similar to previously published work investigating ripples in MTL and cortex^47–50^ to demonstrate the presence of oscillatory events within the ripple band (80 – 180 Hz) comparable to ones previously reported in non-human primates^51^. After identifying candidate events, we further filtered events with a high signal-to-noise ratio (see Figure S8 and Methods) to avoid issues with artefacts or noisily defined events^52^. This procedure revealed candidate MTL-like sharp wave ripples (Figure 5A, n = 4162, median duration = 103 ± 42.08SD msec; see also Figure S8-9). Crucially, despite our wide frequency range, the ripples were localised to a narrow band spanning 90-140 Hz (Figure 5B, bottom). This concurs with previous findings^24,48,51,53^ and suggests these ripple events were generated by a specialised circuit rather than reflecting spurious high-frequency noise spanning the entire frequency range. The simultaneously recorded vmPFC neurons were found to elevate their firing rate around the ripple window (Figure 5C) – and more on trials with ripples (Figure S10A) – which coincided with a ripple sharp wave component (Figure 5D). While ripples have been previously described in human cortex and rodent mPFC, this demonstrates the first evidence of ripples in non-human primate vmPFC within a value-based decision-making context^48^.

**Figure 5:**
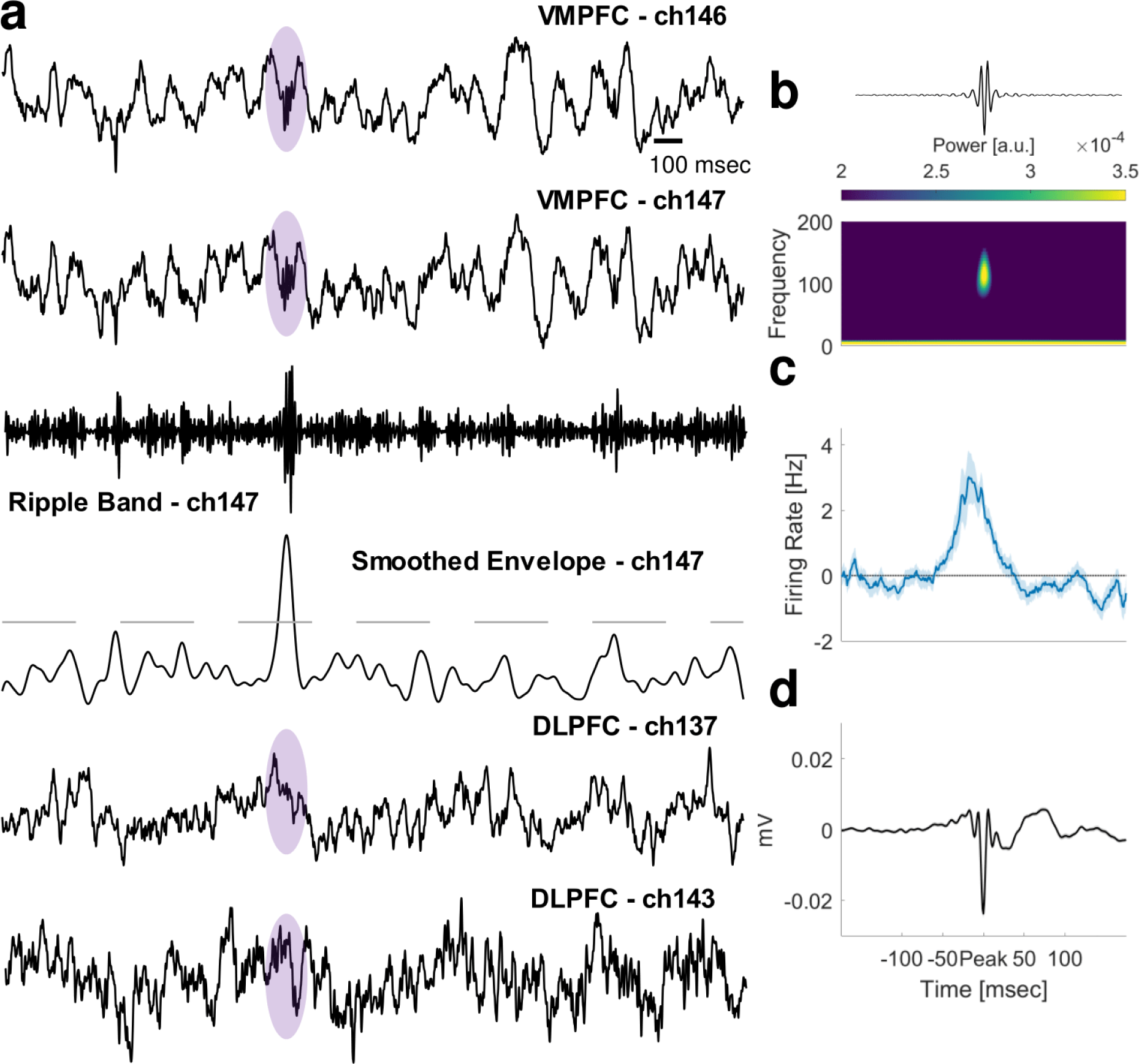
Sharp wave ripples in the vmPFC. **a)** Four simultaneously recorded channels. The top two rows show the wideband LFP signal where a candidate ripple was detected on both electrodes in vmPFC (purple shading). The third row from the top shows the top signal filtered to the ripple band (80 – 180Hz). The fourth row shows the smoothed envelope where the z-score crosses the threshold used to select candidate ripple events. The bottom two rows show the wideband LFP signal of two other channels recorded simultaneously at identical time points. Note that all simultaneously recorded LFP channels are from independent microelectrodes spaced a minimum of 1mm apart**. b**) Top panel: mean oscillatory activity across all ripple band events of all channels together. Bottom panel: average power spectrum of the above events superimposed across the same time period. **c)** The ripple events coincided with increased firing in vmPFC neurons. Error bar represents SEM across vmPFC neurons. **d)** Average high-pass filtered signal highlighting the sharp wave component.

### vmPFC ripples are modulated by value-guided choice

Having identified the presence of ripples in vmPFC, we next asked whether these were modulated by value-guided choice. We localized ripple events in time relative to task events and compared the ripple event frequency in vmPFC with that of the other brain regions. We observed a temporal specificity of ripple events whereby they mostly occurred shortly before and after choice, and at feedback, and they occurred at a much higher frequency in vmPFC compared to the other brain regions (Figure 6A). Furthermore, the periods where ripples in vmPFC occurred most did not coincide with periods of a general tonic increase in the firing rate of the vmPFC neural population, which might have suggested that the recorded events in the previous Figure Simply indexed a tonic increase in the firing of vmPFC neurons, such as what occurs in the other brain regions (Figure S10B).

**Figure 6:**
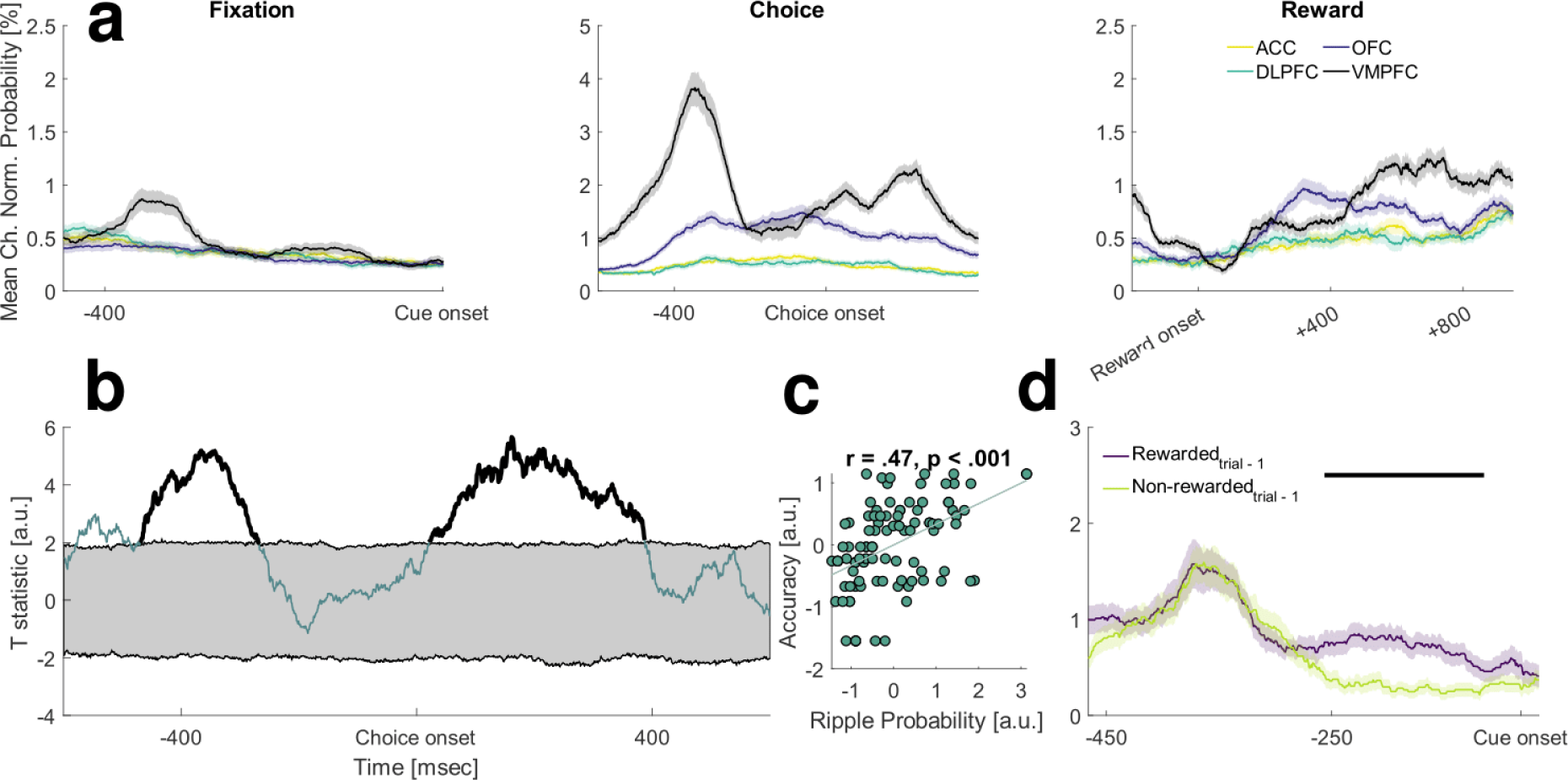
Accuracy and reward modulate ripple proportion during choice and rest. **a)** Average probability of detecting ripples on a trial, averaged across all channels in all four brain regions. Error bars represent SEM across channels within each brain region (ACC = 149, dlPFC = 104, OFC = 133). **b)** Session-level accuracy can be predicted from ripple probability across channels in vmPFC before and after choice. The highlighted black lines represent clusters exceeding the 97.5^th^ percentile of a length-corrected null distribution. The gray shading represents the permutation thresholds of the null distribution. **c)** Correlation taken at peak of b) before choice, each dot represents a vmPFC channel. **d)** When subjects were previously rewarded, more ripples occur in the subsequent fixation period. Error bars represent SEM across vmPFC channels (n = 46). The black bar denotes significance in cluster-based permutation testing (exceeding the 97.5th percentile of the length-corrected null distribution).

Because ripples have been previously associated with task performance (e.g. memory retrieval)^48,49^, we predicted the frequency of these events would be related to behavioural performance in the context of choice accuracy, considering their potential role in model-based planning^17,18^. When we tested this, we observed that ripple event frequency before choice predicted session accuracy (Figure 6BC), meaning that subjects performed better on the sessions where more ripples were observed across channels. This result was not explained by average session reaction time, stimulus set familiarity, or subject identity, as these were regressed out before performing this analysis.

Recent work has shown that theta stimulation during the fixation period disrupts learning in an RL task and this is thought to be due to impaired communication between PFC (OFC) and hippocampus^44^. If ripples play a comparable communication-related role in our task, we should find increased ripple frequency in the fixation period after subjects make correct choices or receive reward. Because the probability of detecting these events on channels was lower in fixation compared to during choice, we performed a median split on channels, picking ones where more ripples were detected, to improve our signal-to-noise ratio. We then tested our prediction and found a higher proportion of ripple events occurring on trials after subjects were rewarded for their choice (Figure 6D); similar to work showing ripple event modulation by previous reward^54^.

Therefore, in addition to demonstrating the presence of ripples in vmPFC, we show these are modulated by task events during value-based choice, and that vmPFC ripples correlate with task performance and reward. This suggests vmPFC ripples may be involved in value-based learning and decision-making processes and offers a further parallel to the well-established planning and spatial navigation processes of the medial temporal lobe.

## Discussion

We investigated neural representations that facilitate flexible, value-guided choice in prefrontal cortex, by taking inspiration from the well-characterised map-like representations in medial temporal lobe and spatial cognition. We demonstrated – for the first time – a grid-like code of value space at choice in the LFP of two separate datasets, showed the same grid-like code in neurons, and demonstrated how such encoding in neurons may be theta phase-dependent. Furthermore, we found grid orientations realign across stimuli sets. Finally, we found sharp wave ripples in the vmPFC of non-human primates and that they are modulated by task events in a value-guided choice task.

The grid-like code was found at choice and represented a navigation angle that embedded both currently relevant choice options as locations in a value space. That this navigation angle revealed a grid-like code suggests the latter is traversed during choice, like in spatial navigation^14,38^. This is important because this navigation angle, together with the length of the vector between the options, contains all the information necessary to determine the correct option choice (decision variable) in such a value space. For example, anything in the upper right quadrant of a value map (Figure 2A) is more valuable compared to anything in the bottom left. Hence spatial navigation problems and making optimal choices in value-guided decision-making tasks may be solved using common neural mechanisms, such as vector-based navigation.

Furthermore, we demonstrate that when subjects must make choices between options which themselves have to first be composed with distinct visual cues, the attributes indexed by those visual cues, such as probability and magnitude, are abstracted away from the sensory properties of the cues to form a latent cognitive map for value. This abstraction is crucial because such a representation of the task structure could enable generalisation across sensory instantiations of this task, or tasks similar to it^16,21^. Notably, this result also corroborates a similar recent fMRI finding where a latent grid-like code of social information (competence by popularity) was found in human vmPFC^13^.

Next, we showed grid orientation realignment across stimuli sets. This demonstrates a non-spatial analogue of the grid realignment observed in spatial navigation when subjects are transferred between – and generalise knowledge across – spatial environments^42^. In addition to deepening the parallel between spatial and non-spatial accounts of grid-like coding, this result suggests the mechanisms underpinning spatial generalisation may also mediate non-spatial generalisation. Furthermore, by showing further parallels, this result provides additional evidence that signals indexed by the hexadirectional analysis, are, indeed, grid cells – an assumption underlying this analysis approach which has not been proven yet.

These results suggest the strong anatomical links between vmPFC and MTL may be important in terms of how representations in vmPFC are encoded and what kind of neural computations we should expect to find in this region^28^. Thus, they may highlight further representations and computations we could expect to find in vmPFC and how these may be implemented. First, like neurons in the hippocampal formation, vmPFC neurons were modulated by theta frequency. This mimics findings from another brain region with strong functional and anatomical links to the hippocampal formation - non-human primate OFC^28,43,44^. Second, like in the MTL, we found oscillatory activity fitting the definition of sharp wave ripples^24,47,48^.

Our data provide the first evidence for a grid-like code of value space in neurons generally, and vmPFC neurons specifically. While this signal was observed to occur when neurons fired the most (theta trough), it is not clear from our data whether theta phase-dependent encoding of information is a ubiquitous property of vmPFC. One argument to support that claim is finding a chosen value signal on a separate theta trough, following the one with a grid-like code, in two separate but related measures. When this is taken together with the finding that vmPFC neurons contributing most to the grid-like code are an independent population from the one contributing most to the chosen value code, it suggests separate neuronal populations in vmPFC encode different task variables sequentially (or in parallel) and in a theta phase-dependent manner. If theta phase-dependent encoding is a ubiquitous property of vmPFC neurons – as our data suggest – then this may be an explanation for the scarcity of vmPFC neuronal findings: if longer firing rate time windows are averaged over time, as is common in the field, phase-dependent signals may average out. A related question is whether the strength of value-related signals would more generally improve if they were to be assessed against the underlying oscillatory phases in which they may be embedded in^44^.

Finally, we detected ripple events, another common feature found in MTL, predominately in vmPFC, and to a much lesser degree in the other three regions. This corroborates rodent work finding ripple events in mPFC^24^ and previous non-human primate work in relation to the frequency band within which they were found^51^. It remains to be seen whether these ripples originate in the hippocampal formation or whether they are generated within vmPFC itself. If the latter, the synchronisation of ripples between the two regions would be interesting to observe^24^. Despite showing ripple events in vmPFC were modulated by task events (e.g. when they occurred relative to choice), and that ripple event probability predicts choice accuracy, it is not clear what their precise role is in such tasks. Work within MTL suggests a role in the retrieval and consolidation of information^41,48,49,55^, in the replay of task states and choice trajectories^17,56^, or in model-based planning^18^. Note that in our task, subjects needed to compositionally combine cues into options to make choices with continuously novel stimulus sets (Figure 1A). The relationship between ripple probability and choice accuracy or reward delivery in the previous trial implies these patterns may be a mechanism by which state value associations may be consolidated or reinforced. Exploring these possibilities are exciting avenues for future work.

## Conclusion

We demonstrate the existence of a cognitive map for value-guided choice in vmPFC LFP theta frequency and vmPFC neurons revealed by the navigation angle of currently relevant choice options. Together with previous non-human primate fMRI work showing a cognitive map in a value space during passive observation of options across trials^12^, these two studies jointly speak towards a new representational architecture for value during value-guided choice. Within this perspective, vmPFC represents economic decision-making tasks in a way that is neither predicted nor explained by pre-existing theories of value-guided choice^1,2,5,57,58^. Furthermore, it means making a choice between two options cannot be accounted for only through a comparison of two or more options in a common currency format^1,5,57^. Rather, these results suggest that a cognitive map is constructed in the vmPFC along the dimensions that are relevant for choice, with this representation being employed when a choice needs to be made between options along those dimensions. This is particularly useful when value needs to be inferred based on having structural knowledge of a task, e.g.^12^, or constructed on the fly in novel contexts. Taken together, our results suggest that the well-described neural mechanisms in MTL which support cognitive maps during spatial navigation have been redeployed by vmPFC for building cognitive maps of conceptual knowledge.

## Acknowledgements

We would like to thank Dan Bush for very helpful discussions. S.V. was supported by the Leverhulme Doctoral Training Programme for the Ecological Study of the Brain. E.G. was supported by MRC grant MR/N013867/1. T.E.J.B. was supported by a Wellcome Trust Senior Research Fellowship (grant no. 104765/Z/14/Z), Wellcome Trust Principal Research Fellowship (grant no. 219525/Z/19/Z) together with funding from the James S. McDonnell Foundation (grant no. JSMF220020372). L.T.H. was supported by a Henry Wellcome Fellowship (098830/Z/12/Z) and Henry Dale Fellowship (208789/Z/17/Z) from the Wellcome Trust; a NARSAD Young Investigator Grant from the Brain and Behavior Research Foundation; and the NIHR Oxford Health Biomedical Research Centre. S.W.K. was supported by the National Institute for Mental Health (grant no. F32MH081521) and the Wellcome Trust Investigator Awards (nos. 096689/Z/11/Z and 220296/Z/20/Z). The funders had no role in study design, data collection and analysis, decision to publish or preparation of the manuscript.

## Author contributions

S.V., T.H.M., E.G., J.L.B., S.W.K. conceived the study. E.G., L.T.H., and S.W.K. collected the data. S.V., T.H.M., E.G., T.E.J.B., J.L.B., S.W.K. analyzed the data. All authors interpreted the data. S.V., T.H.M., J.L.B., and S.W.K. wrote the paper with input from all the authors. T.H.M., J.L.B., and S.W.K. supervised the project.

## Methods

### Dataset 1: Subjects & Recordings

Subjects were two adult male rhesus monkeys (M. mulatta), subjects M and F. Both weighed 7-10kg and were ∼4 years of age at the start of this experiment. Their daily fluid intake was regulated to maintain motivation in the task. All experimental procedures were approved by the UCL Local Ethical Procedures Committee and the UK Home Office and carried out in accordance with the UK Animals (Scientific Procedures act). For additional details on subjects, electrophysiological methods and recording locations of Dataset 1, see^1,2^

We recorded neurons in ACC (dorsal bank of anterior cingulate sulcus, area 24c), dlPFC (both dorsal and ventral banks of sulcus principalis, areas 9/46), OFC (primarily in the medial orbital gyrus, area 11/13), and vmPFC (medial to the medial orbital sulcus and within the rectus gyrus, area 14). Recordings were done with single microelectrodes spaced apart at a minimum distance of 1 mm. For a detailed description of the recording protocol and recording locations, see^1,2^. In total, 724 neurons (198 – ACC, 156 – DLPFC, 195 – OFC, 160 – vmPFC) and 497 LFP channels (149 – ACC, 104 – DLPFC, 133 – OFC, 97 – vmPFC) were analysed.

### Dataset 1: Task

Subjects performed a two-alternative forced choice task where both options were simultaneously presented, one on each side of the screen (Figure 1A). Each option was parametrized by two attributes, reward magnitude and reward probability. Each attribute had five reward levels, yielding a total of 10 possible cues (images). Magnitude cues represented juice amount in arbitrary units (0.15, 0.35, 0.55, 0.75, 0.95). Probability cues represented the probability of receiving the reward (10%, 30%, 50%, 70%, 90%). Each attribute could appear either on top or the bottom of the screen. The presentation of the cues and the position of attributes with respect to the screen were sampled uniformly to avoid any biases in the stimuli presentation. On each trial, the subject therefore saw four cues appear on the screen, two on the left side indicating the magnitude and probability of the left option and two on the right side indicating the magnitude and probability of the right option. These four cues appeared simultaneously to the subjects, and they were free to saccade to sample information about each option before making a choice. Note that this is a separate dataset from ref^1^ with the same subjects. Subjects chose by moving the joystick in the direction of the option they were choosing. Following the cue presentation and subjects’ choice, a short pre-feedback period followed, where subjects saw the selected option but did not receive reward. This was followed by the feedback period where they received the outcome for the selected option followed by a fixation cross. A stimulus set was repeated for three to four sessions.

Following this, subjects underwent a learning task in which they learned the mapping between ten novel visual cues and their corresponding reward magnitude and reward probability levels. In the learning task, subjects only learned the mapping between visual cues and their corresponding reward magnitude/probability level by observing a secondary reinforcer (filled bar), where the filled height of the bar indicated the level of value to assign to the novel stimulus (see ref^3^ for learning task details). In the learning task, subjects never made choices between options where an ‘option’ would span two dimensions. Therefore, all choices in the two-alternative forced choice task presented here were novel as subjects never had to combine individual attributes (reward magnitude and reward probability) into an option and use that information to make choices. For additional details on the task see^2^.

### Dataset 2: Subjects & Recordings

The second dataset is from an ongoing project where a full dataset has been collected in one subject. The subject was one adult male rhesus monkey (M. mulatta), subject A. The subjects weighed ∼14kg and was ∼11 years of age at the start of this experiment. The subject was implanted with a titanium head positioner for restraint. On the basis of preoperative 3T MRI and stereotactic measurements, a recording chamber was subsequently implanted on the left hemisphere and with its centre 26mm along the anterior-posterior (AP) coordinate plane. Postoperatively, gadolinium-attenuated MRI imaging and electrophysiological mapping of gyri and sulci confirmed chamber placement. Craniotomies were then performed inside the chamber to allow for neuronal recordings. During each recording session, up to 4 multi-channel linear electrodes (26-32 channel V-PROBES, Plexon, Dallas, TX) were lowered into the brain through a grid with custom-built manual microdrives to measure neuronal activity. A Plexon Omniplex system was used to record neuronal data at 40 kHz and local field potential data at 1kHz. Single-unit isolation was achieved with manual spike sorting with a Plexon Offline Sorter. Neurons were randomly-sampled, with no attempt to pre-screen for responsiveness. We recorded neuronal data from vmPFC, defined as all sites medial to the medial orbital sulcus and within the rectus gyrus (area 14) spanning AP 35-42mm. Recordings were also made in OFC defined as sites between the medial and lateral sulci (areas 11/13) spanning AP 35-42mm. In total 252 channels from vmPFC and 209 channels from OFC were analysed.

### Dataset 2: Task

Subjects performed a two-alternative forced choice task with sequential presentation of two options. Each option was parametrized by two attributes depicted by cues (images): reward magnitude and reward probability. Each attribute had ten reward levels, yielding a total of 20 possible cues. Magnitude cues represented juice amount in arbitrary units (0.17, 0.34, 0.51, 0.68, 0.85, 1.02, 1.19, 1.36, 1.53, 1.70). Probability cues represented the probability of receiving the reward (9.1%, 18.2%, 27.3%, 36.4%, 45.5%, 54.6%, 63.7%, 72.8%, 81.9%, 91%). The first option (‘Option 1’) was presented for 700 msec on either the left or right side of the screen and the subject was allowed to freely fixate the option. Both attributes of the option were presented simultaneously, and the position of reward magnitude and reward probability were randomized across trials. The second option (‘Option 2’) was presented for 700 msec and the subject was allowed to freely fixate the option. If Option 1 appeared on the left side of the screen, Option 2 would appear on the right side of the screen. The order of option presentation was randomized across trials. Once central fixation was re-acquired, both options were presented for choice. Choice was required through fixation for the chosen option for 500ms. The chosen option was then presented paired with reward or no reward according to its associated magnitude and probability. Feedback on correct/incorrect choice was provided via a change in background colour.

### Behavioural analysis

We computed behavioural accuracy on a per-session level by computing whether the subjects would pick the option with higher expected value. Expected value (EV) was defined as:

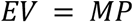

Where M refers to the magnitude of reward an option yields and P refers to the probability of obtaining that reward if an option is selected. For trial-wise analyses, correct trials were defined as trials where subjects chose the option with the higher expected value. The psychometric curve was computed by collapsing choice behaviour across all recorded sessions and computing the probability of selecting the right option as a function of the rank *EV* difference between both options.

### Neuronal CPD analysis

The spiking information of each neuron was aligned on a trial-by-trial basis according to the subjects’ choice onset and smoothed with a 100-millisecond boxcar filter incremented by 10 milliseconds. We used the coefficient of partial determination (CPD):

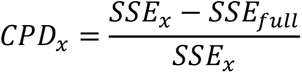

to investigate whether a canonical representation of value exists in the four brain regions that we recorded from. In the equation above *SSE*_*x*_refers to the sum of squared errors in the regression model excluding the regressor of interest *x*, *SSE*_*full*_refers to the sum of squared errors of the full model and *CPD*_*x*_, therefore, refers to the CPD of regressor *x*. As is visible from the equation, the CPD for a regressor tells us how much a regressor alters the sum of squared errors of a dependent variable. For our analyses, the CPD provided information about how much variance the EV of the chosen option (chosen value) and the EV difference between the chosen and unchosen option (chosen value difference) explain for each neuron.

GLM 1 had two regressors:

1. Intercept (constant term)
2. Chosen value (continuous value)

GLM 2 had two regressors:

1. Intercept (constant term)
2. Chosen value difference (continuous value)

The chosen value regressor was defined by:

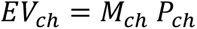

Where EV is the expected value of the chosen option, *M*_*ch*_ denotes the magnitude and *P*_*ch*_ denotes the probability of the chosen option. Similarly, the chosen value difference regressor was determined by subtracting the above-chosen value regressor from an equivalent regressor created for the unchosen option. The CPD was estimated at each timepoint for each neuron within a region and the average across all neurons per region was plotted over time. Furthermore, the CPD values were averaged within a larger 300 millisecond time window locked to choice onset, corresponding to conventional time windows used previously to identify value-related representations emerging in these four regions^1,4^

### Determining the statistical significance of the CPD

We determined the significance of the chosen value and chosen value difference in individual regions by permutation testing. To do this, we created a null distribution. The null distribution was created by decoupling the dependent variable (firing rate) from the independent (regressor) variable. This was achieved using a shuffling procedure repeated 1000 times. For each of the 1000 samples, we generated a mean null CPD per region by averaging across all neurons within one brain region. This resulted in a null distribution for each region. We then computed the mean and standard deviation of the CPD of the null distribution and converted the mean empirical CPD into a z-score:

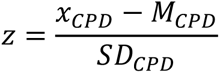

Where *z* refers to the z-score. The z-score of each region was converted to a p-value by a Fisher z transformation. *x*_*CPD*_ is the mean empirical CPD for a given region, while *M*_*CPD*_and *SD*_*CPD*_are the mean and standard deviation of the CPD of the null distribution, respectively. A region would have been considered statistically significant if the region’s p-value were below the statistical threshold of p < .05. All regions except vmPFC in Figure 1 were significant at a threshold of p < .001.

### Local field potential preprocessing

Local field potential (LFP) data was sampled at 1000Hz. We cleaned and pre-processed the data using established methods by removing line noise and harmonics using the discrete Fourier filter from the Fieldtrip toolbox^5^. Following visual inspection of the pre-processed wideband LFP activity, we observed noise artefacts in the data. To remove them, we rejected trials if the wideband LFP amplitude on a given trial around the subjects’ choice (50 milliseconds before choice to 350 milliseconds after choice) exceeded 3SD of the mean. This removed 0-3% trials per channel.

### Value GLM used for regressing out value

The GLM used for regressing out value had the following regressors in the design matrix:

1. Intercept (constant term)
2. Chosen value difference (continuous value)
3. Trial difficulty (continuous value; absolute value difference)
4. Chosen magnitude (rank value from 1 to 5)
5. Chosen probability (rank value from 1 to 5)
6. Unchosen magnitude (rank value from 1 to 5)
7. Unchosen probability (rank value from 1 to 5)
8. Attribute position (binary regressor, top or bottom)
9. Chosen side (binary regressor, left or right)
10. Reaction time (continuous value)

The trial difficulty regressor was used as a side-agnostic value regressor (i.e. whether subjects’ chose left or right) and was defined as the absolute value difference between the expected value on the left and right side.

### Hexadirectional analysis

To find evidence of a grid-like code, we used the hexadirectional or quadrature filter analysis by closely following the approach taken in previous work^6–9^. Specifically, we used the method reported previously for LFP data in^9^. Briefly summarized, for each session we first obtained an angle estimate which was derived as follows: A value space was defined on a Cartesian coordinate system where reward magnitude spanned the *x* axis and the reward probability spanned the *y* axis. The left and right option subjects were making choices between were embedded as locations in this two-dimensional space. That is, each option represented a location in a reward magnitude by reward probability value space. Based on such an embedding, we were able to compute an angle between the left and right choice options by using the *x* axis as the base of a hypothetical triangle. The angle on each trial was computed by:

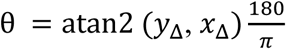

Where *y*_Δ_ denotes the vector length difference of the reward probability and *x*_Δ_denotes the vector length difference of the reward magnitude. Centring the analysis on the angle difference between the left and the right option allowed for unbiased sampling across the angle space compared to selecting the angle difference between the chosen and unchosen option. For each session, we then sorted trials by the obtained angle. This was done because the grid orientation was estimated in a cross-validated way where two-thirds of the data were used to estimate the grid orientation for each channel using a GLM:

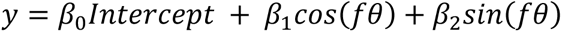

Where *f* denotes the tested symmetry spanning from 4 to 8 folds and *θ* denotes the angle. The obtained betas from the *sin* and *cos* component of the regression equation were then converted into a grid orientation by:

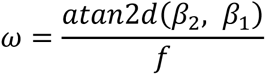

This grid orientation was used to adjust the angle of the held-out third of the data by subtracting the grid orientation from the angle and generating an adjusted angle for each trial (*θ*_*adjusted*_ = *θ* − *ω*). These adjusted angles were used in a GLM on the held-out third of the data where a cosine regressor was used to test for a grid-like representation of value:

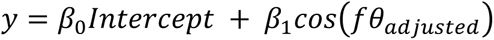

This analysis was performed on theta band (5-8Hz) LFP activity like in previous work ^39^ which was transformed using a Morlet wavelet using the *cwt* function implemented in Matlab R2020a. The data was then down-sampled to 100Hz to speed up analyses. The obtained GLM procedure computed using this data was repeated three times (i.e. for all three folds) generating three independent regression betas per channel. The whole procedure was repeated for all recorded channels within each of the four recorded regions. This procedure was first computed on neural activity averaged within the original 300 millisecond time window, like the analysis estimating the significance of CPD values. Furthermore, we computed the strength of the grid-like code for value across a longer time window centred on the choice epoch when subjects made choices.

In Dataset 2, the same approach was used to find evidence of a grid-like code. Due to the sequential nature of Dataset 2, angles were computed between Option 1 and Option 2 which was not possible in Dataset 1 where both options were presented simultaneously. Similar to Dataset 1, the strength of grid-like coding was estimated in a time window before choice – we used a time window from 100 to 500 msec following the presentation of Option 2, which was used in previous work showing the emergence of value signals in prefrontal cortex during this time window^1^.

### Determining significance of hexadirectional analysis

For each channel, we first averaged the three independent regression betas that were obtained from the hexadirectional analysis. This yielded channel-level beta estimates which indicated the strength of a grid-like code on a per-channel basis. To avoid concerns about whether regression estimates from individual LFP channels can be statistically treated as independent units of analysis, we further averaged channel estimates within a brain region to obtain independent session-level estimates. This yielded session-level beta estimates for each brain region. To perform statistical inference on these quantities, we employed two separate methods. First, to determine a significant population-level effect across sessions, we performed one-sample t-tests across sessions for each symmetry to investigate whether the beta estimates were significantly different from zero. The crucial prediction is that these betas would yield a positive, significant *t*-value for sixfold symmetry but not control symmetries. That would imply a higher theta band LFP signal on angles aligned with the grid axis compared to angles that were misaligned with the grid axis. Second, to estimate the significance of the effect on a per-session basis, we compared session-level beta estimates to a null distribution by shuffling the dependent variable 1000 times on a per-session basis before running the hexadirectional analysis. The session-level betas were considered significant if they exceeded the 99^th^ percentile of the null distribution. These analyses were always performed for control symmetries as well. Similar to the first approach described above, session-level significance was estimated based on session-level averages to avoid concerns about whether individual channels can be considered as independent units of analysis.

To further verify the signal was hexadirectionally modulated, we used the computed grid orientation estimates to align and bin the held-out trial angles into aligned and unaligned trials. This was done in a way where the trial-by-trial angles and neural activity were always aligned and subsequently binned with cross-validated grid orientations. For each channel, we therefore obtained a series of six angle bins for both aligned (0, 60, 120, 180, 240, 300) and misaligned (30, 90, 150, 210, 270, 330) trials that spanned the whole value space. We first averaged the trials for each bin of a channel and then averaged all channels within a session, similar to the procedure used to determine the significance of a grid-like code in individual regions.

Across all recording sessions of Dataset 1, there was variance in the number of trials per session, spanning from 151 to 277 trials. Because the hexadirectional analysis is sensitive to angle sampling, we removed six sessions with the lowest number of trials to improve the grid orientation estimates and reliability of our results. This removed sessions with fewer than 200 trials per session (removing 11.27% of recorded channels with the lowest number of trials).

### Hexadirectional analysis control frequencies

To determine whether the grid-like code was present only in theta band (5 – 8Hz) or other frequencies as well, we used a similar approach as in ref^9^ by bandpassing LFP signal into individual frequencies (beta: 12 – 40Hz, low gamma: 41 – 60 Hz, high gamma: 61 – 100Hz), transformed the data using Morlet wavelets within each band, summed the obtained signal within each band, and downsampled the data to 100Hz to speed up analyses. We repeated the hexadirectional analyses described in the previous section on this data.

### Distorting the cognitive map via behavioural model fitting

To test whether choices were biased by a distorted representation of either reward probability or magnitude, we fitted three prospect theory models of increasing complexity on a session-by-session basis, using equation from ref^10^, similar to other recent work^11,12^. At the core of each model were two parameters which measured the distortion of the magnitude and probability axis:

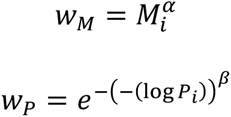

In these equations, *M*_*i*_refers to reward magnitude for state *i* and *⍺* refers to the magnitude distortion parameter. *P*_*i*_ refers to the reward probability for state *j* and *β* refers to the probability distortion parameter. The obtained weights were used to compute subjective expected value:

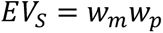

This yielded subjective expected values for the left and right option. We then computed the probability of choice on a trial-by-trial basis using:

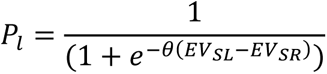

The models were then fitted to minimise *E*:

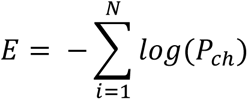

That is, we estimated a set of parameters to minimise the sum of the log-probability of the chosen option (*P*_*ch*_) across all trials (*i* = 1, … *N*) in each session. The optimisation was done in Matlab using the fmincon function. The distortion parameters were initialized at 1 to avoid a prior bias which would favour distortion along either axis. The parameters had an upper bound at 1 as in ref^11^. The choice temperature parameter θ was initialized at the upper bound (set at 10). This means the model initializes the fitting procedure with the assumption of high choice precision and mitigates a scenario where low initial choice temperature (very stochastic decision-making) would potentially falsely lead to larger distortion estimates.

The first extension of this model added a choice noise parameter to account for potential attentional drift across trials:

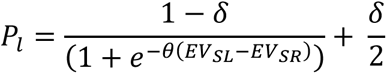

This parameter separately accounts for differences across trials which would not be accounted for by the choice temperature parameter. The parameter was initialized at 0 to avoid the assumption subjects had choice noise due to similar reasoning as the one provided for the choice temperature and was bound at 1.

The final model included a side repetition bias, as defined in ref^12^:

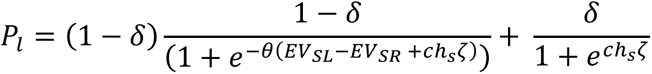

The side repetition bias tests whether a subject is likely to select an option if they have selected that option (*ch*_*S*_) on the last trial (coded as +1 or -1).

All three models were separately fit to subjects’ session-level data and compared using the Bayesian Information Criterion (BIC)^13^. The winning model (model 2) had a BIC score improvement of 8.005 relative to model 1. However, a paired t-test across sessions between model 1 and model 2 showed no statistical significance in model fit (*t*53 = 0.28, *p* = ns). Because we did not have apriori hypotheses about which of these models would best account for subjects’ data we used the one with the best BIC score (model 2).

Using the behavioural magnitude and probability distortion parameters we computed the state-by-state difference in the subjective magnitude and probability estimate compared to a non-distorted case where both parameters are fixed at 1 and indicate no distortion. This was also repeated after computing the theoretical expected value for the whole magnitude by probability value space and comparing it to the subjective expected value for that space. Differences in both measures indicate for which attribute or expected value states can the subjects be said to have over- or underweighted their value.

Using the obtained parameters, we then rescaled the reward magnitude and reward probability levels which were used to compute the navigation angles in the hexadirectional analysis. The remainder of this procedure was identical to the main hexadirectional analysis. This yielded a separate set of channel-level beta estimates indicating the strength of a grid-like code for a distorted representation of the cognitive map which were compared to the original betas reported in the main manuscript with paired t-tests.

### Grid orientation realignment and consistency analysis

A key assumption of hexadirectional analyses is that the signal being indexed is generated due to firing statistics of conjunctive grid cells which fire more when movement is aligned with the main axes of grid firing fields^6^. However, this has so far not been proven. One way in which this claim has been strengthened previously is by showing that grid orientations measured using the hexadirectional analysis approach are stable across days and brain regions^7, 14^. Crucially, a computational property of grid cells is that their grid orientations realign when moved between environments. In other words, the measured grid orientation from one environment may change when the grid orientation from the same grid cell is measured in another environment^42^. We were able to test this prediction due to our task design where the same environment (i.e. a set of five reward magnitude and reward probabilities) was continuously assigned to new stimuli sets (i.e. different sensory observations). This allowed for a different type of cross-validation from the one presented in Figure 2. Namely, in line with grid orientation alignment, we expected grid orientations to remain stable when cross-validated across sessions within the same stimulus set but realign across sessions of different stimulus sets^15^. To first validate that reasoning, we estimated the grid orientation for each channel using the method described in Hexadirectional analysis and then computed pairwise distances between each channel using the equation:

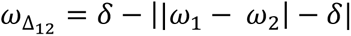

Where *δ* is a constant denoting the width of the grid field for a given symmetry level which was set to 60, *ω*_1_ is the grid orientation from one channel and *ω*_2_ the grid orientation for another channel. This yielded a channel-by-channel distance matrix from which only the upper triangle was used for analyses due to the matrix being symmetric. From this matrix, we computed the average distance for each session for vmPFC channels as a function of whether the distance was obtained from other sessions with the same stimulus set or with another stimulus set. Crucially, in our analysis, we did not use within stimuli set distances that came from the same session as this would potentially artificially inflate how small within stimuli set distances are due to signal correlations that may exist across channels within a session (i.e. it may have increased our false positive rate). In the final step, we created two averages, one coming from the same stimuli set and another one coming from different stimuli sets. This distance was compared to the null distribution where we shuffled the label at the previous step 1000 times (same stimuli set, different stimuli set) and used a Fisher Z transformation to assess statistical significance. The subsequent analysis where we estimated the grid consistency followed an identical procedure to the one described in Hexadirectional Analysis. As we observed a weaker grid-like code on the first day a stimulus set was presented, we excluded channels from such days in this analysis to ensure avoiding false negatives that could have occurred due to badly estimated grid orientations.

### Neuronal phase-aligning, grid-code, and chosen value analyses

To estimate phase-dependent firing in neurons, we first estimated theta phase in the same frequency range in which we found the grid-like signal (5 – 8Hz) on every trial. On every trial, we then binned theta phase into 10-non overlapping phase bins within which the raw neuronal firing rates were binned. For each phase bin, we slid a larger 300-millisecond window over time and averaged the neuronal firing rates within this window for each neuron and trial at each window increment. This allowed us to estimate neuronal firing rates in a theta phase-specific manner, estimate how theta phase dependency may change over time, and crucially, allow capturing more than one theta cycle within the larger 300-millisecond time window which increased the robustness of the phase-dependent firing. Figure 4A shows the phase-dependent average across vmPFC neurons collapsed across the choice epoch while Figure 4B shows the phase-dependent average of an example neuron without collapsing across the choice epoch. To determine whether neurons exhibited phase-dependent firing we first mean-subtracted each neuron’s activity at individual phase bins using the average across all phase bins and examined the effect of phase in a one-way ANOVA. In the supplementary materials, we performed further post-hoc paired t-tests across neurons where we compared the neuronal firing across neurons at each phase bin. The post-hoc paired t-tests across neurons were used to determine the ‘Preferred’ and ‘Non-Preferred’ phases that were used to illustrate the phase-dependent firing in Figure 4C; the ‘Preferred’ phase corresponded to the phase bin with the largest relative neuronal firing activity across neurons compared to other phase bins while the ‘Non-Preferred’ phase corresponded to the phase bin with the lowest relative neuronal firing activity across neurons.

We compared the relationship between neurons and oscillatory data in three different ways. First, we bandpass filtered our LFP activity across all vmPFC channels within the theta range in which we found the grid-like encoding of value (5 – 8Hz). We then averaged each channel across trials and further averaged the data across all channels. Through this, we obtained the average theta oscillatory activity across all channels shown in Figure 4D which enabled us to define theta-based seed windows (labelled with numbers from 1 to 4) within the original 300 millisecond window before subjects initiated their choice from Figure 2B in which we report the grid-like code. Each seed window was centred on the maximum (peak: 1, 3) and minimum (trough: 2, 4) of the average theta oscillatory activity in 60 msec bins. This bin length allowed us to minimize the amount of temporal overlap, i.e. it allowed us to use two non-overlapping windows (peak, trough) of an 8Hz oscillation (1 cycle = 125 msec) which was the highest frequency tested and reported in the main text. These seed windows were used to compute the grid orientations of vmPFC channels which were then used to compute the strength of grid-like encoding in vmPFC neurons. Therefore, the only difference between these grid orientations was the part of the average theta oscillatory cycle within vmPFC they occurred in. For each seed window, all control symmetries were computed.

Next, instead of relying on the average theta oscillatory cycle, we phase-aligned neuronal firing rates in time on a trial-by-trial basis within the original time window. For each channel, we computed the theta phase for every trial and used the temporal differences in phase peaks across trials to fully align the neuronal firing rates in vmPFC neurons identified on those channels. After alignment, the neuronal firing rates of each neuron were smoothed with a 25 msec boxcar filter to minimize the possibility of contaminating phase-specific signals occurring across cycles. We repeated this across all channels and their corresponding neurons. The strength of this approach is that it allows us to examine the properties of neuronal firing rates fully aligned to theta oscillatory cycles. The limitation of this approach is that it jitters the precise information of each time point with respect to task epochs (e.g. Choice onset). Due to this, we computed a histogram of the average times that were used to time-align these theta cycles (i.e. where peaks occurred) to temporally localize the time points with respect to task epochs (Figure S7G). This showed a correspondence between this approach and the above ‘seed window’ approach. As in the approach above, we used the grid orientations of vmPFC channels to compute the strength of grid-like encoding in vmPFC neurons. This allowed us to hold grid orientations constant and examine the emergence of a grid-like code in neuronal firing rates. We determined whether a grid-like code in neuronal firing rates exists by estimating the hexadirectional beta for each neuron and doing a one-sample t-test against zero across all neurons. If there is a grid-like code in neurons, this test should yield a significant positive t-value. To determine whether such significant positive t-values could be obtained by chance we used cluster-based permutation testing. For each neuron, we randomly shuffled the firing rates across trials 1000 times – using identical shuffles across time to preserve temporal smoothness - before computing the betas at each time point. We then performed t-tests across neurons for each shuffle and compared the empirically obtained t-values against the null distribution (97.5th percentile of a length-corrected null distribution).

To examine the firing properties of neurons with a high grid-like code, we averaged the neuronal firing within the window with a significant grid-like code in Figure 4G of for two neurons. The averaged firing rates were z-scored across all trials. The firing rates were then further averaged as a function of each state presentation (i.e. each presentation of reward magnitude and reward probability). The presentations of each state were averaged for the left option and right option separately. The obtained rate map for each neuron and each option was then convolved with a 3×3 Gaussian kernel with sigma = 0.5.

The phase-aligned neuronal firing rates were also used to examine the emergence of a chosen value signal using a CPD approach like the one described in Neuronal CPD analyses. To determine the significance of the chosen value CPD, we used the same length-corrected null distribution permutation testing approach as described in this section.

### Sharp wave ripple detection

To detect sharp wave ripples, we pre-processed the raw LFP signal by removing line noise and band-passing it between 1 and 250Hz. This band-passed signal was further filtered to the ripple band (80 – 180Hz) as in previous non-human primate work^16^. To detect candidate ripple events, we used ripple detectors like previous work^16–18^. In brief, the ripple band-filtered LFP signal was smoothed with a 4SD Gaussian. The envelope of the smoothed trace was then computed by using the Hilbert transform. This envelope was z-scored. On the z-scored envelope we looked for events with a total length of at least 25 msec, of which at least 15 msec needed to exceed 3SD throughout their duration. If several events were recorded where peak to peak was separated by less than 40 msec, these were combined into one ripple event. These candidate events were then further processed by a non-negative matrix factorisation with the aim of improving our signal-to-noise ratio and selecting events that have clearer spectral properties^16^ due to recent concerns with respect to the signal-to-noise ratio of ripple estimation^19^. We subjected the wideband spectral signal of each ripple event to a non-negative matrix factorisation with three clusters and 10 replicates. We then proceeded to compute for each of the three obtained clusters their low-rank approximations of the original spectral signal. For each cluster, we then computed the estimated signal-to-noise ratio for each frequency (by dividing the spectral signal by the summed squared error the projection of each cluster gave rise to relative to the original spectral signal). We then defined a minimum signal-to-noise threshold that was used to discard events below this threshold. We used a signal-to-noise ratio of 5 compared to the original 3 used by^16^. The events passing this criterion needed to further pass the final criteria by which we selected only events as ripples when the projection of the cluster with the highest signal-to-noise ratio had a clear unimodal peak that was within the ripple band, as defined in ref^16^. Crucially, this allowed for ripple events where the projection of a cluster with high signal-to-noise ratio can also be observed in other frequencies (e.g. due to theta activity).

To determine whether the observed ripples were associated with increased oscillatory power in the ripple range, we computed scalograms using bandpass-filtered and line noise-removed LFP data using Morlet wavelets in the frequency range of 1 and 200 Hz. These were always centred on the peak ripple signal using the ripple detector. We computed the power by squaring the magnitude and then performed baseline decibel normalisation with respect to the first 300 msec of the fixation period. Similarly, we computed the peak-centred firing rate of all neurons recorded within vmPFC across all channels by concatenating all trials where ripples occurred for each neuron. This was followed by computing averages on a per-neuron basis and smoothing the raw data with a 50 msec boxcar filter. Finally, we baseline subtracted the firing rates using activity of the first 300 msec on the same trials. The ripple band oscillation and sharp wave components were averaged in the same manner as described above. The ripple band oscillation was bandpass-filtered to the frequency described in the manuscript while the sharp wave component oscillation was low-pass-filtered as in ref^16^.

### Ripple analyses

To determine the relationship between ripple frequency and choice accuracy we computed subjects’ choice accuracy on a session-by-session basis and examined whether it is related to the probability of detecting ripples on LFP channels across sessions. To do this, we used a GLM where we predicted session-level choice accuracy using channel-level ripple frequency. Crucially, this GLM contained co-regressors for subject identity, session-level reaction time, and the number of days a stimulus set was presented for to rule out confounding explanations that could have driven choice accuracy across sessions. To determine the significance of the relationship between ripple probability and choice accuracy we asked whether the obtained t values from the GLM exceed the 97.5th percentile of a length-corrected null distribution.

To determine the relationship between rewarded and unrewarded trials in relation to ripple presence, we split trials according to whether they were rewarded and unrewarded on trial *t* and averaged the ripple rasters on the subsequent fixation/rest period which occurred before the next trial started. To determine significance, we used cluster-based permutation testing where we shuffled trials and investigated whether ripples survived the 97.5^th^ percentile of the length-corrected null distribution.

## Supplementary Figures

**Figure S1.**
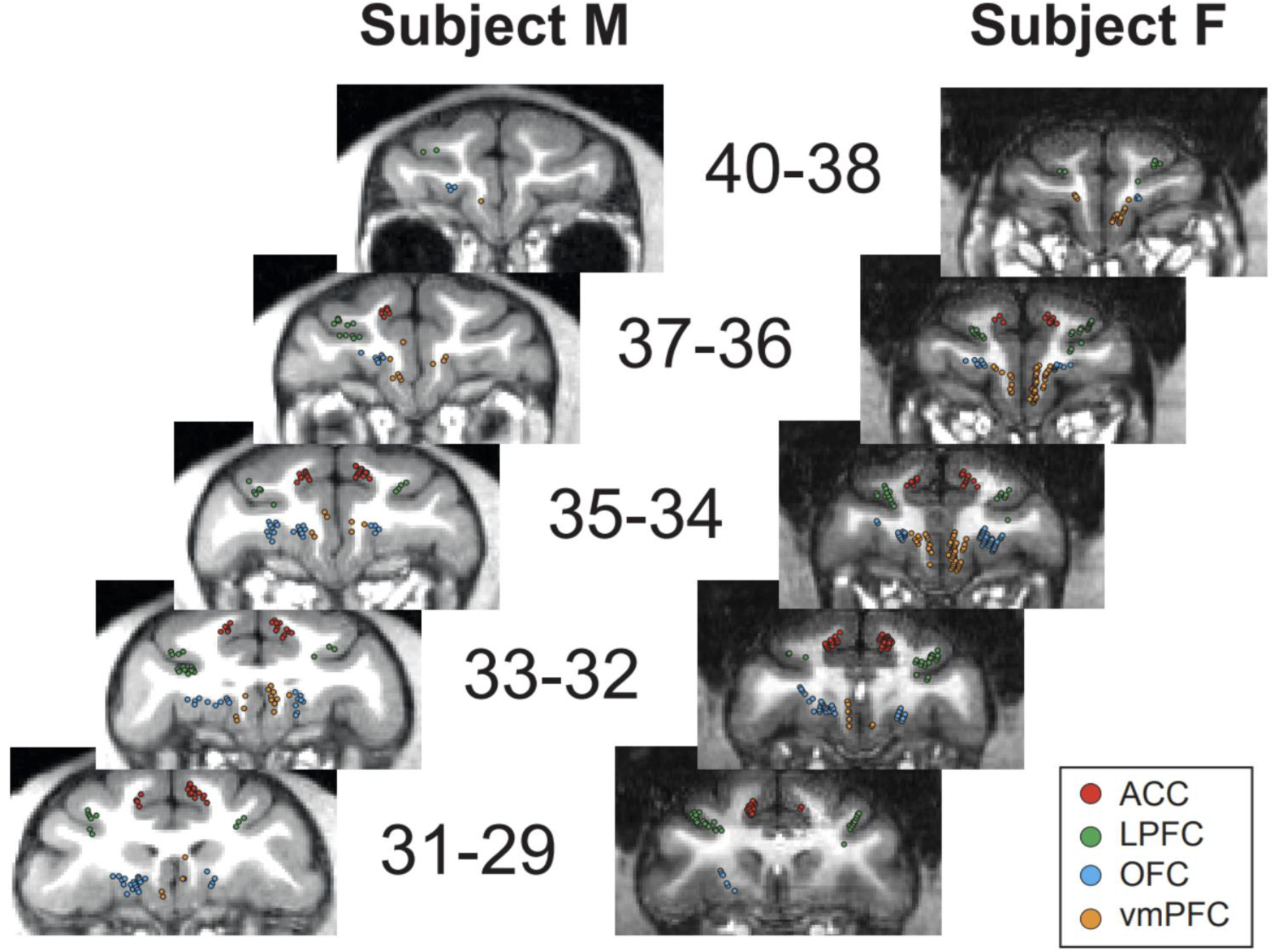
Neuronal recording locations. Taken from^2^. Images are each monkey’s MRI at various coronal planes. Numbers signify the distance in millimetres from anterior posterior 0. Each dot represents on electrode from one recording location.

**Figure S2.**
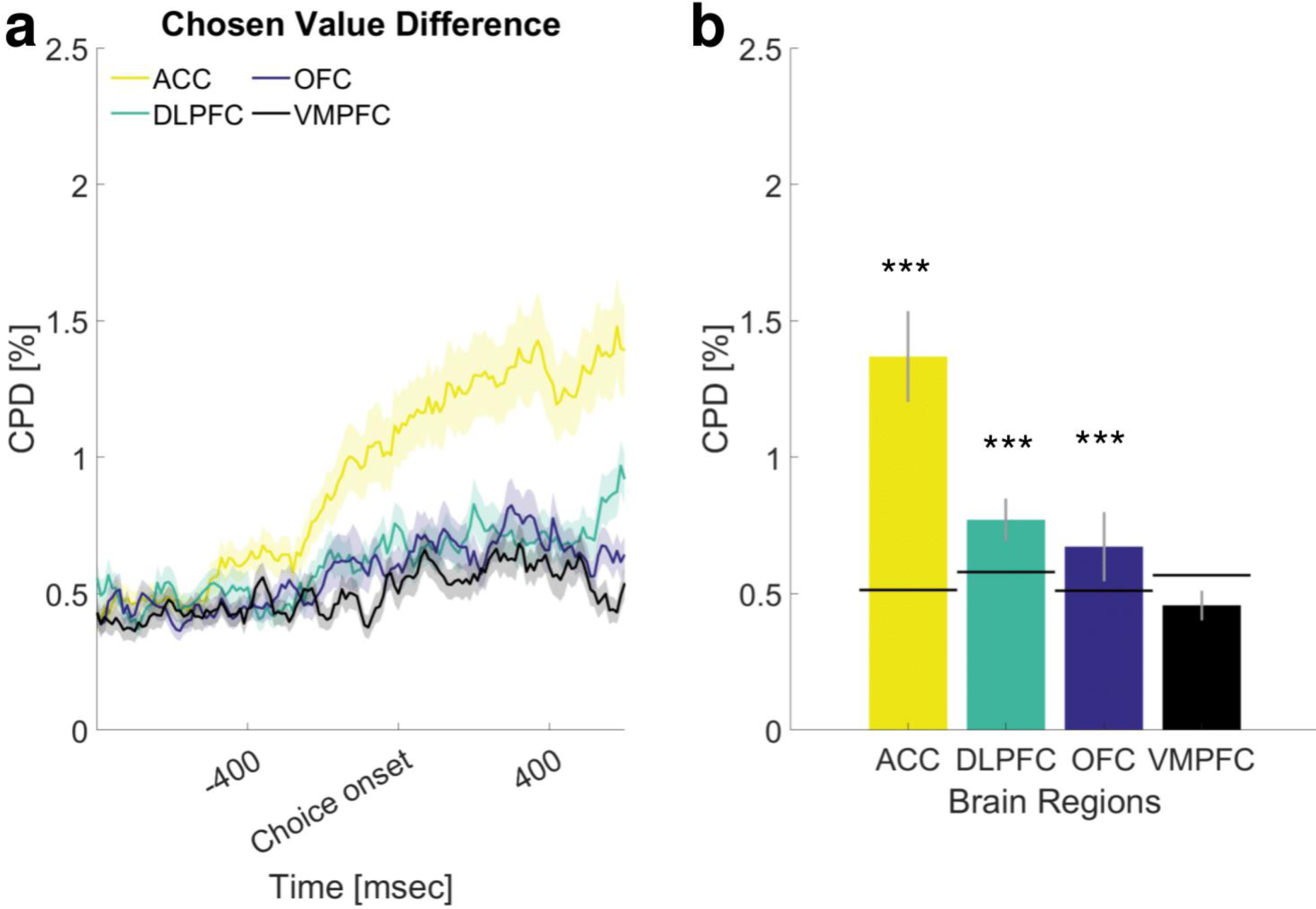
Chosen value difference. **a)** Chosen value difference coefficient of partial determination (CPD) for individual brain regions across the choice epoch. The thick line represents the mean response, the shaded area represents the standard error of the mean (SEM) averaged across neurons within brain region. Cue onset was approximately 560 msec before choice onset (mean reaction time across sessions). **b)** Mean CPD for the same regressor as in a) for individual brain regions within a 300-millisecond time window before subjects initiated their choice. The black line denotes the 95th percentile of a null distribution for visual purposes, obtained through permutation testing (1000 permutations). *** p < .001. Error bars represent SEM across neurons within the brain region.

**Figure S3.**
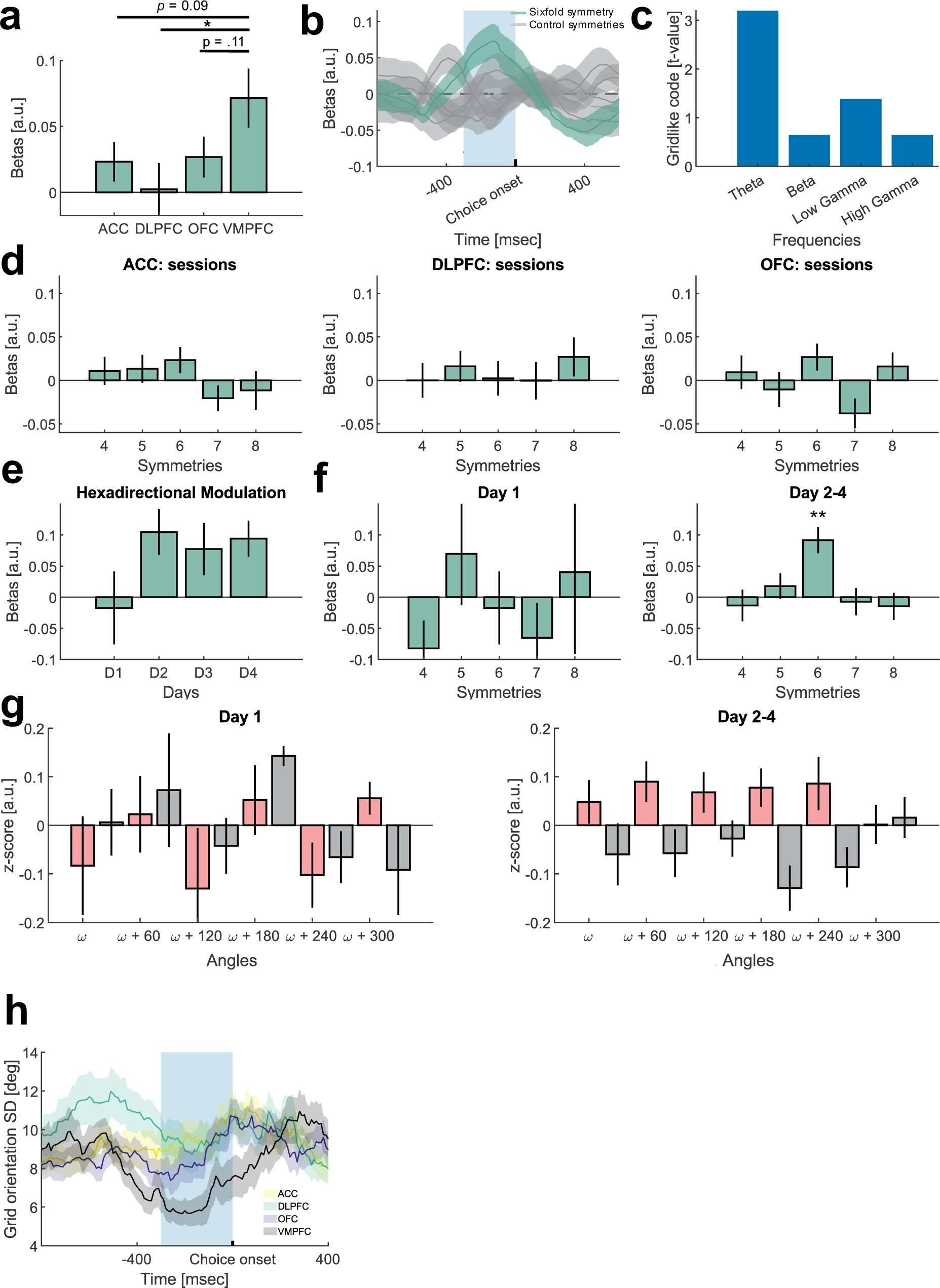
Grid-like code across regions, frequencies, and across stimulus set repetitions. **a)** Hexadirectional (sixfold) modulation across the four recorded regions. The height of each bar represents the average across sessions for each brain region. Note that the vmPFC bar is identical to the one from Figure 2B. Error bars represent SEM across sessions for each brain region. * p < .05. **b)** Hexadirectional (sixfold) modulation in vmPFC for sixfold (green) and control (4-8) symmetries (gray). Blue shading denotes the 300-millisecond window before subjects initiated their choice. The thick gray and green lines denote session averages of betas obtained from the regression with which we determined the existence of the grid-like code. Shaded areas denote SEM across sessions. **c)** Grid-like code in vmPFC across frequencies defined based on ref^9^. We find no significant grid-like code in control frequencies. Notably other work has reported a grid-like code in gamma^20^. Note the t-statistic for Theta is identical to the one reported for Figure 2B. **d)** The effect reported in Figure 2B for other regions. Height of bars and SEM conventions are identical to Figure 2B. **e)** The Hexadirectional (sixfold) modulation effect reported in Figure 2B split as a function of the number of days of stimulus set exposure (a stimulus set was swapped every few days). There was a trending effect indicating the grid-like code became stronger on subsequent days of a stimulus set presentation compared to the first day it was presented (t14 = 2.12, p = 0.052). **f)** The hexadirectional (sixfold) modulation effect reported in Figure 2B shown for Day 1 and the average of Day 2-4. ** pBonferroni < .01. **g)** Sixfold periodicity reported in Figure 2D shown for Day 1 and the average of Day 2-4. **h)** Decrease in grid orientation variance in vmPFC compared to other brain regions. The average standard deviation in the grid orientation across the three folds significantly decreased in vmPFC relative to other regions during the period in which we observed the grid-like encoding of value (ACC: t55 = 2.59, p = 0.01, dlPFC: t46 = 2.42, p = 0.02, OFC: t48 = 2.35, p = 0.02).

## Distorting the cognitive map for value

The analyses presented in Figure 2 make a key assumption: a fully veridical representation of the cognitive map for value. That is, the hexadirectional analysis used to test for grid-like encoding assumes that the representational geometry for neighbouring magnitude or probability levels corresponds to how the experimenter defines those quantities. However, human and non-human primates exhibit choice biases^12,21,22^, with this being especially well studied in the human decision-making literature based on prospect theory^21^.

If the cognitive map contains representational distortions, as opposed to being represented veridically, then those representational distortions may be related to distortions in behaviour. We tested this idea by first fitting different prospect theory models as in ref^11^ using equations from ref^10^ on a session-by-session basis. The prospect theory model with the lowest BIC indicated an overweighting of low magnitude and probability values and underweighting of high magnitude and probability values across recording sessions relative to not being distorted (Figure S4A).

Next, we asked whether behavioural distortions are related to neural distortions (i.e. a distorted representation of the cognitive map). Because previous work reported grid-like representations in ACC, OFC, and vmPFC^7,23^, and vmPFC was significantly stronger only compared to dlPFC but not the other two regions (Figure S3A), we initially collapsed our analysis across these regions on a per-session basis. We then asked if distorting the magnitude and probability values using the best-fitting behavioural parameters would improve the grid-like code in these regions. We found that distorting both magnitude and probability according to behaviour (Figure S4B) resulted in a reduction of the grid-like code. This effect was driven specifically by vmPFC (Figure S4C, *t*14 = 2.30, *p* < .05). It is important to highlight that this effect did not replicate in the second independent dataset (Figure S5). It should, therefore, be treated as preliminary and warrants further investigation. Different possible factors may have led to a lack of replication: e.g. lower statistical power in Dataset 2 (fewer total sessions), differences in task design, and or differences in training regimes.

If the result is replicated in future work, it suggests vmPFC contains a veridical representation of the cognitive map, while other regions downstream in the choice process may distort this veridical representation. While the exact relationship between neural and behavioural distortions of this kind is currently unclear, this approach provides a way of bridging biases from rational choice theory and distortions in the representational geometry of the structure of value-based decision-making tasks.

**Figure S4.**
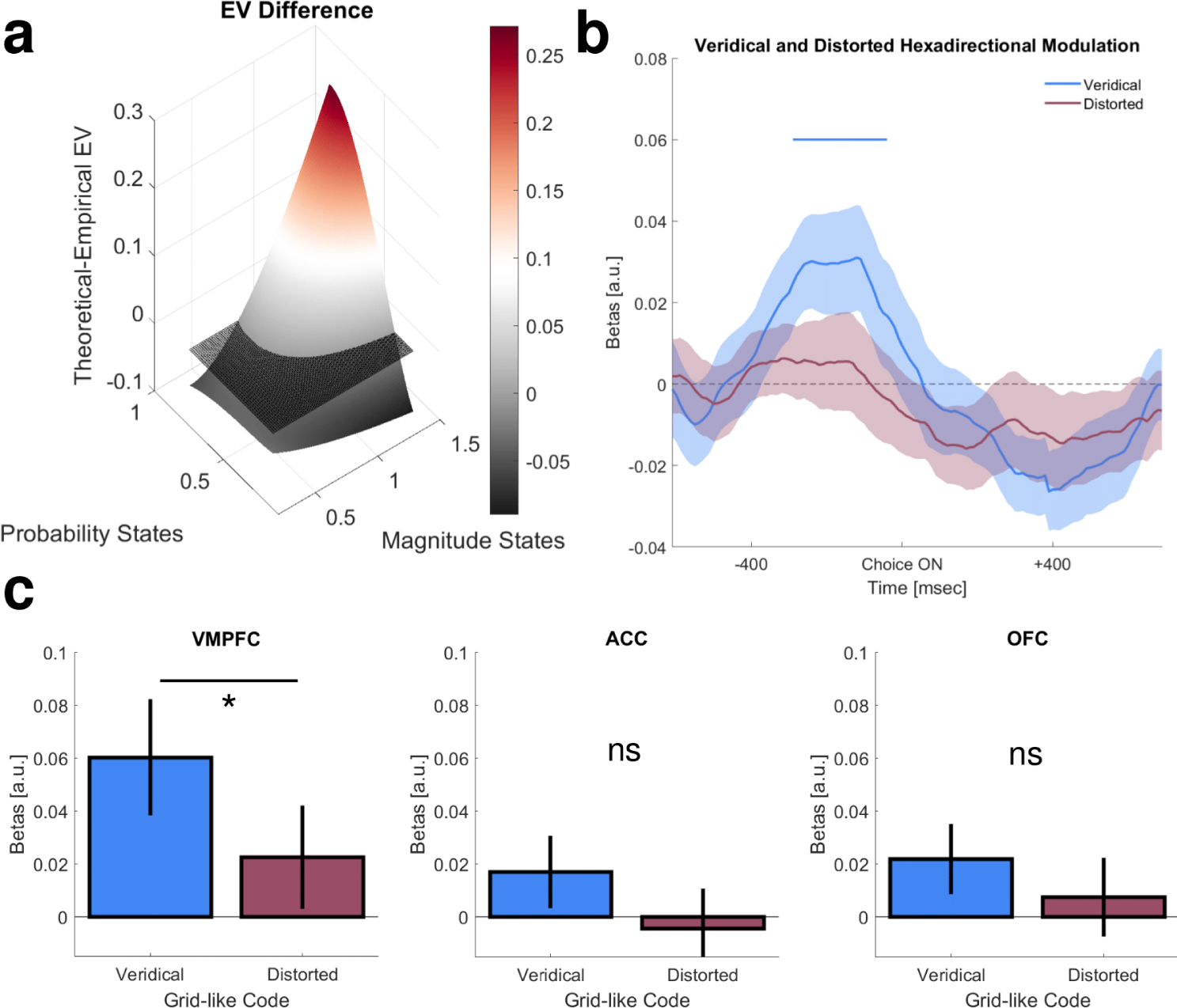
Distorting the cognitive map. **a)** Difference in expected value (EV) between the theoretical and empirical EV space. The theoretical EV space has no distortions in either magnitude or probability, while the empirical EV space has distortions for both magnitude and probability. The coloured plane shows the difference between the theoretical – empirical EV across sessions. High values (red) indicate behavioural underweighting of option combinations with high EV. Low values (black) indicate behavioural underweighting of option combinations with low EV. The flat stripped plane is used for visual purposes to indicate no difference between the theoretical and empirical EV, **b)** a comparison of the average hexadirectional effect across vmPFC, ACC, and OFC for the veridical (blue) cognitive map (analogous to averaging Figure 2C across the three regions) and the distorted (red) cognitive map. The thick blue line corresponds to a significant difference between the veridical and distorted conditions obtained from cluster-based permutation testing (exceeds the 97.5^th^ percentile of a length-corrected null distribution). The error bars represent SEM averages across sessions of vmPFC, ACC, and OFC. **c)** Comparison of the veridical and distorted hexadirectional effect across the three regions within the time window of Figure 2B. Error bars represent SEM across sessions within brain regions. * p < .05.

**Figure S5.**
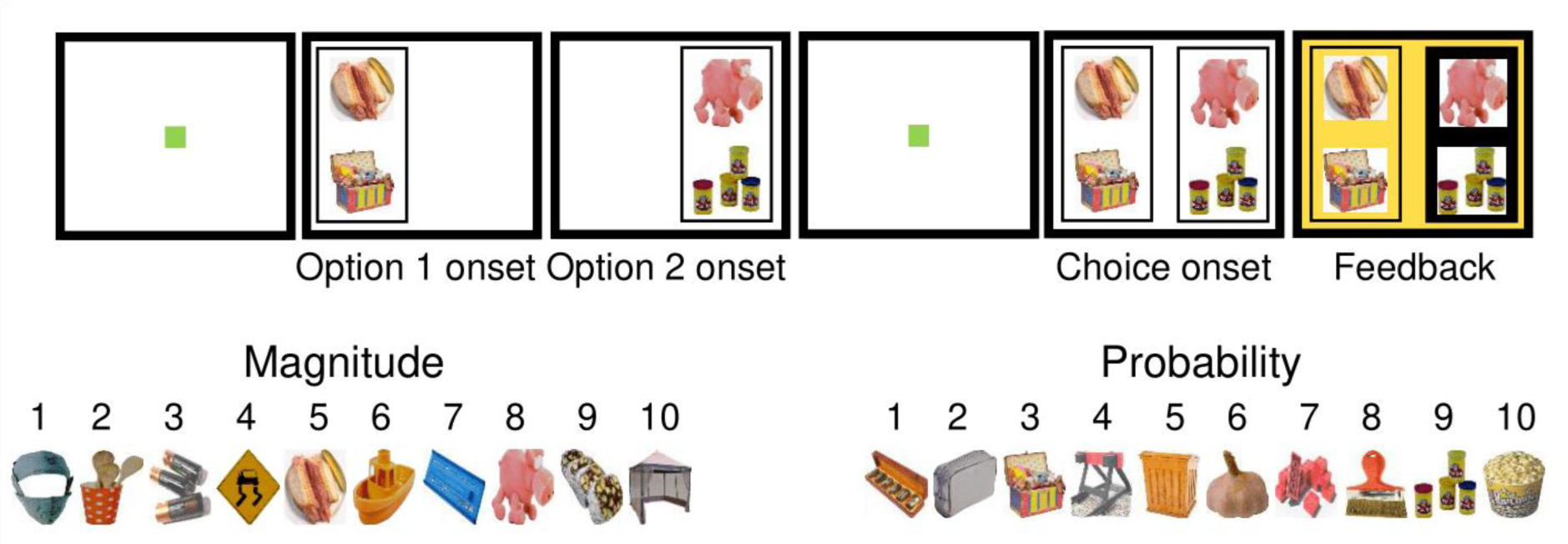
Independent dataset task. Two options were presented sequentially for 700ms (Option 1 onset, Option 2 onset). The subject was allowed to freely fixate each of them. Once central fixation was re-acquired (after Option 2 onset), both options were presented simultaneously for choice (Choice onset). A choice was made via fixation of the chosen option for 500ms. The chosen option was rewarded according to the corresponding magnitude and probability (Feedback). Choice feedback was provided through a change in background colour.

**Figure S6.**
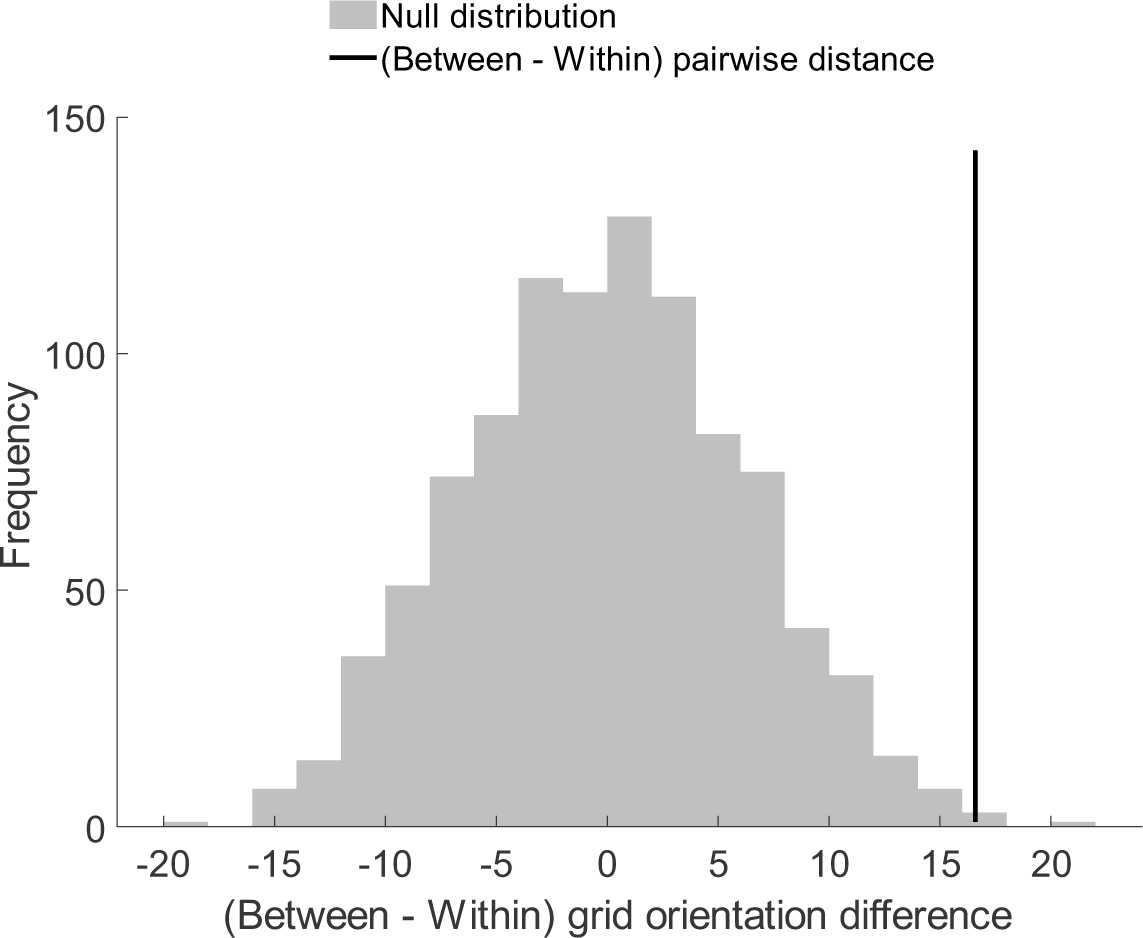
Grid orientation control analysis. Grid orientation distance realignment is significant when controlling for the distance of pairwise comparisons. This control analysis aims at controlling for a possible temporal drift of the grid orientation due to time passing across sessions or other confounds such as training time. This control analysis repeats the grid orientation realignment analysis such that the time differences (i.e. degree of exposure to a stimulus set) of pairwise comparisons, happening either within- or between-stimuli sets were held constant. While this reduced the sample size – because data from vmPFC was not always sampled across individual stimulus set repetition levels due to the difficulty of recording from this region – it replicates the effect presented in the main manuscript. The colour convention is identical to Figure 3E.

**Figure S7.**
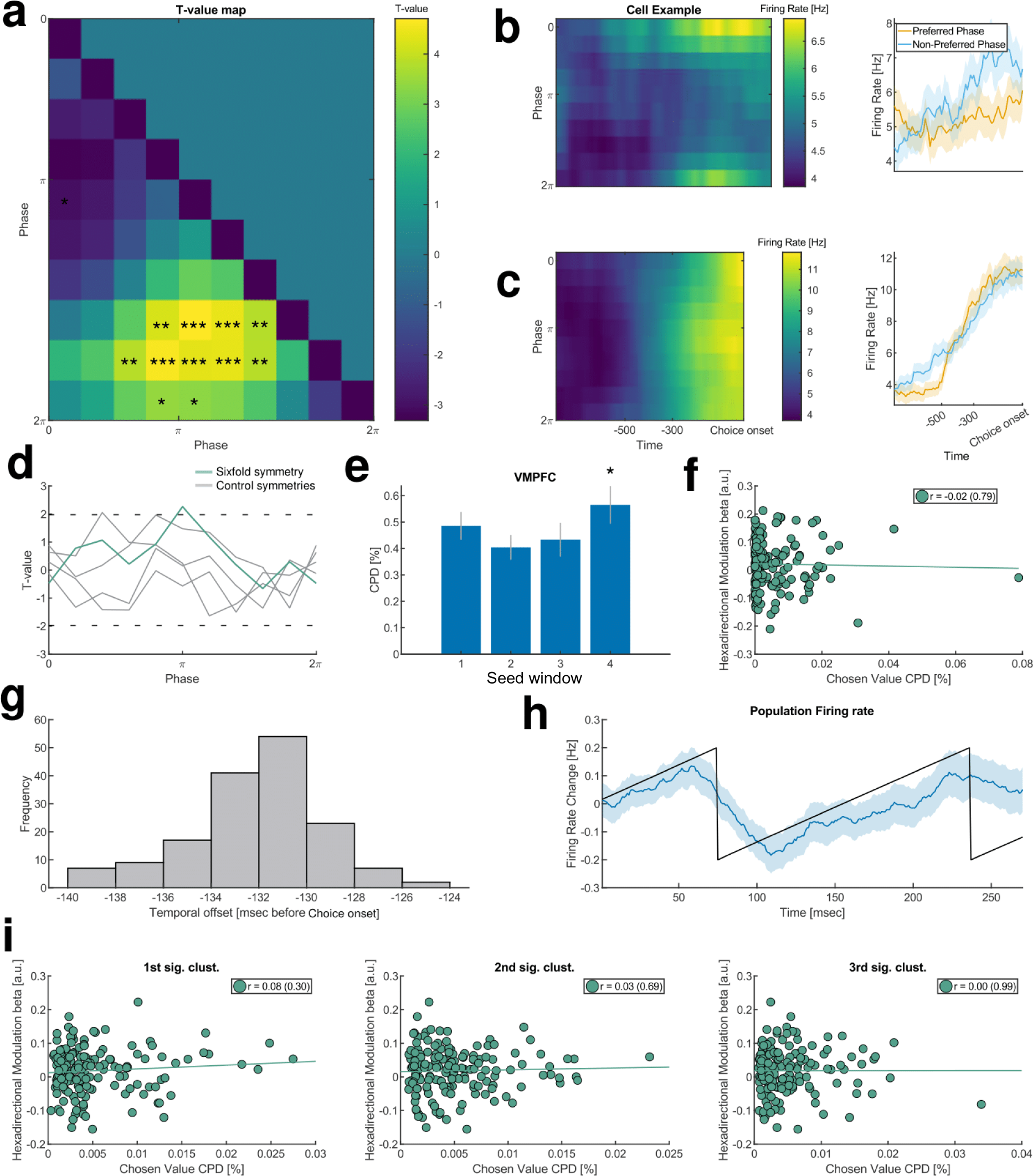
Firing rate properties of vmPFC neurons and additional control tests for a grid-code in neurons. **a)** a phase-by-phase t-value map showing theta frequency phases where vmPFC neurons have significantly higher / lower firing rates. This map was obtained by performing t-tests on firing rates for every pairwise phase comparison. For example, the first column across all rows indicates neurons fired less at theta phase peak (indicated by 0) as theta phase progressed towards the trough (indicated by π). **b)** firing activity of an example neuron which exhibited a firing pattern opposite to that of the majority cells – increasing firing most at theta peak instead of trough. Preferred phase refers to a portion of the theta phase where cells, on average exhibited the highest firing activity. Non-Preferred phase refers to a portion of the theta phase where cells, on average, exhibited the lowest firing activity. Error bars represent SEM across trials for that neuron. **c)** firing activity of an example neuron showing no discernible modulation by phase. **d)** A grid-like code can be found in vmPFC neurons when firing rates are averaged across the same phase of all theta cycles before choice within the 300-millisecond window before subjects initiated their choice from Figure 2B. Error bars represent SEM across vmPFC neurons. Dotted black lines correspond to significance thresholds based on one sample t-tests. **e)** Chosen value CPD before choice. The numbering on the x-axis follows the convention from Figure 4D. Significance was obtained through permutation testing (n = 1000). * p < .05. **f)** The obtained CPD values in the significant seed window (4) from e) do not correlate with vmPFC neuron grid-like code betas. **g)** Temporal localization of the phase-time aligned signal from Figure 4G. Negative values denote milliseconds before subjects made a choice. The peak frequency approximately corresponds to the time of the peak LFP signal. **h)** vmPFC neurons exhibit theta modulation over time, the firing rates are aligned according to one theta cycle and plotted over time, thus keeping the neuronal firing aligned to all phases of theta. Error bars are SEM across vmPFC neurons. **i)** The significant clusters where chosen value was represented in Figure 4L do not correlate with the significant cluster where the grid-like code was observed in Figure 4G.

**Figure S8.**
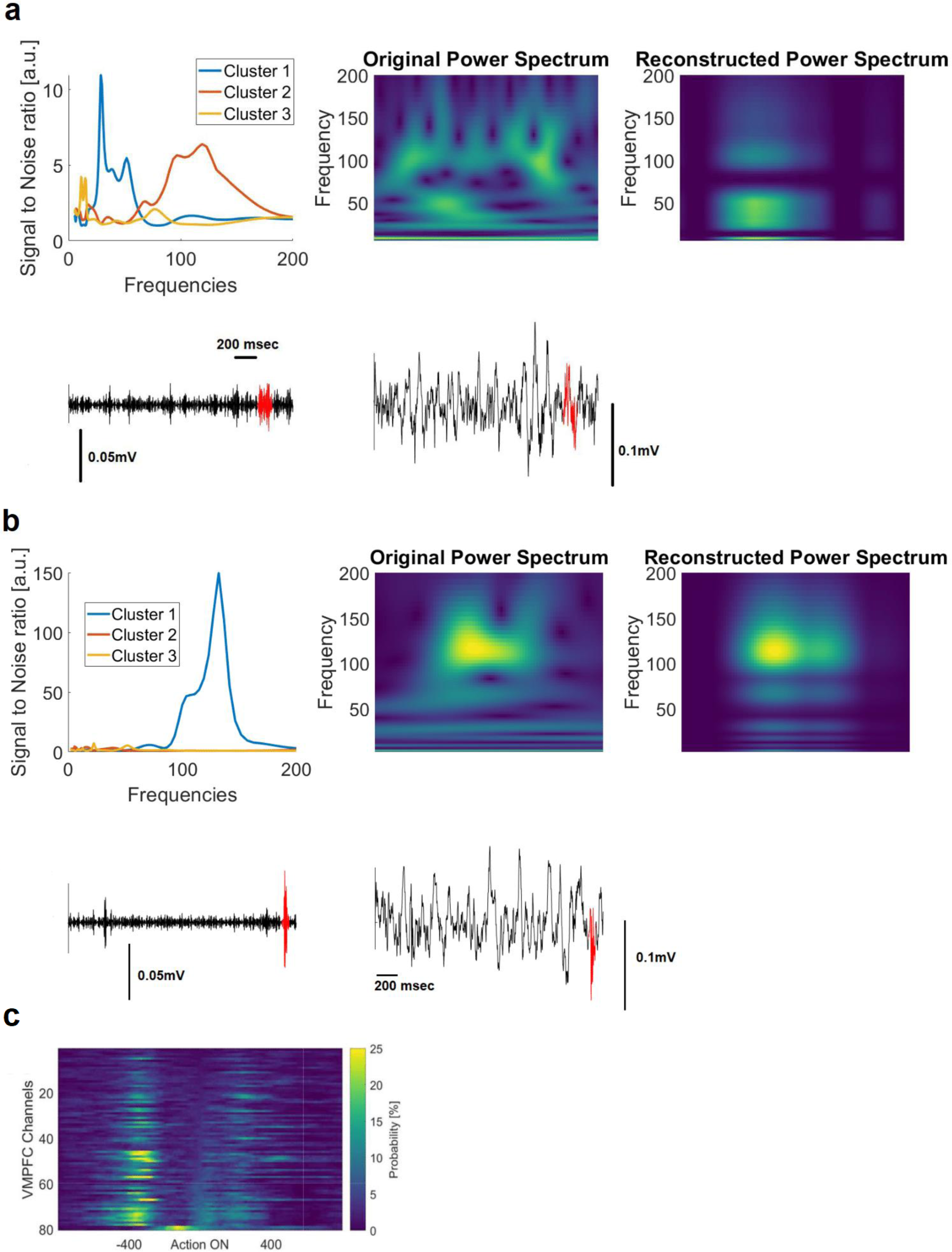
Second-order filtering of candidate events. **a)** Example of a candidate ripple event which was initially detected as a candidate event but was rejected during our second-order filtering procedure. Top left panel: Signal-to-noise ratio for the three identified clusters as a function of the frequency space. Top middle panel: original power spectrum of a candidate ripple event. Top right panel: Reconstruction of the original power spectrum based on the obtained clusters. Bottom left panel: ripple band oscillatory activity (black) with super-imposed candidate ripple events detected by the ripple detector (red). Bottom right panel: wideband oscillatory activity (black) with super-imposed candidate ripple event detected by the ripple detector (red). We computed the power spectrogram (top panel middle) for candidate events (bottom left panel – ripple band activity, bottom right panel – wideband signal). We then performed non-negative matrix factorisation on the power spectrograms with three clusters like ref^51^. For each cluster, we computed its signal-to-noise ratio across frequencies marginalising over time. Ripple events that we used in the analyses presented in the manuscript were events where the principal cluster (cluster with the highest signal-to-noise ratio) exceeded our predefined threshold level and had a clear unimodal peak. Such events could then reconstruct where the dominant spectral signal occurs (e.g. in the ripple frequency domain). **b)**. Example of an accepted event at the second level. The panel conventions are identical to a). **c)** Probability of detecting ripple events across vmPFC channels.

**Figure S9.**
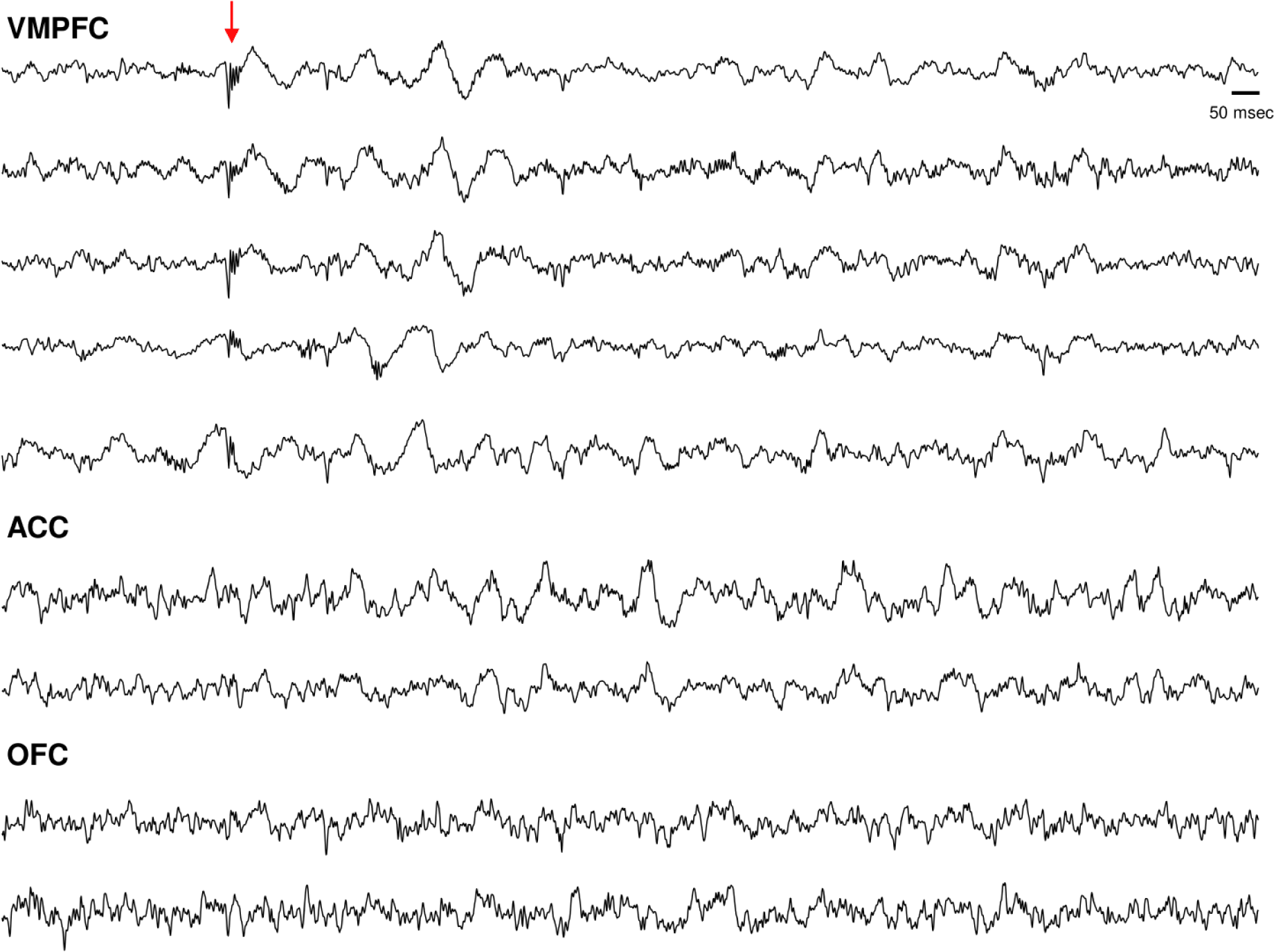
Example session with all recorded channels organised according to brain region. All simultaneously recorded channels from one session at identical timepoints; highlighting cases where events could be detected simultaneously on several channels within vmPFC but not outside vmPFC.

**Figure S10.**
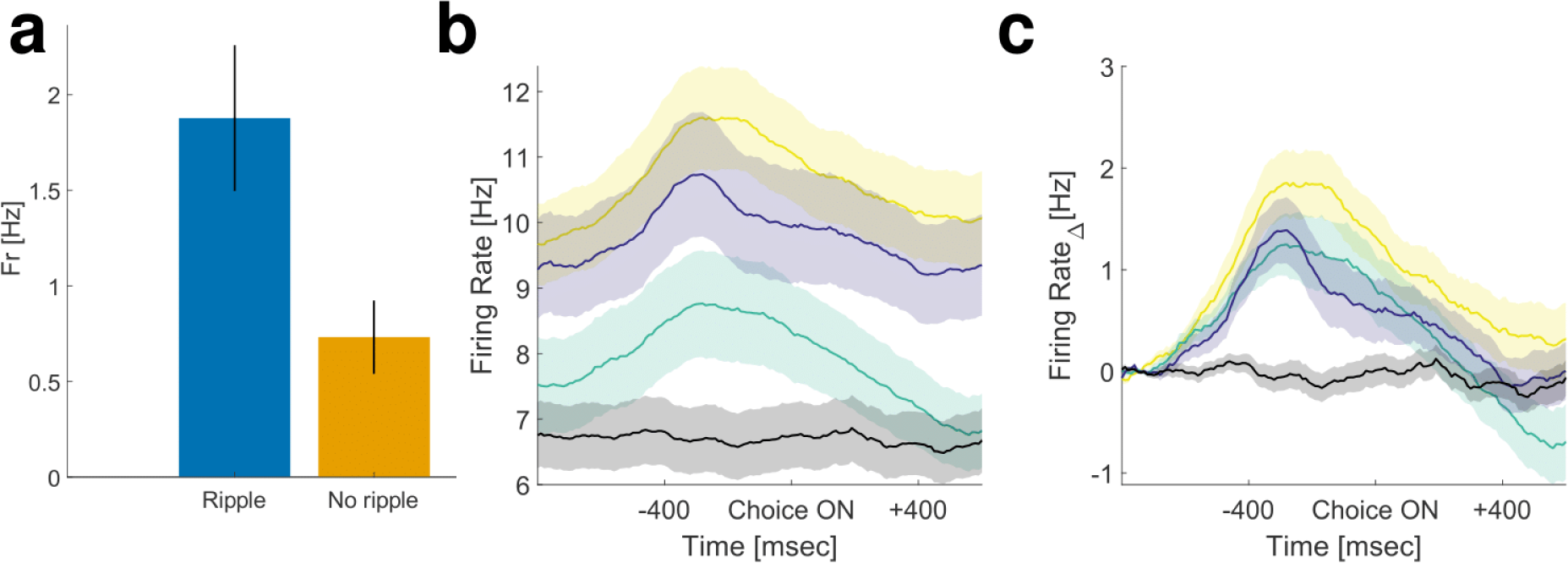
Firing rate on ripple and no ripple trials and firing rate plots over time across regions. **a)** The mean firing rate of vmPFC neurons on ripple trials was higher compared to the firing rate of non-ripple trials (t159 = 3.36, p = 9.67e-04). This was assessed by comparing the average firing rate of vmPFC neurons within a 100 msec window centred on obtained ripple peaks and compared to the firing rate aligned to oscillatory troughs within the same time period but on trials with no ripples detected. Error bars represent SEM across vmPFC neurons. **b)** Mean firing rate before and throughout choice across all regions. Error bars represent SEM across neurons within a brain region. The color convention is identical to Figure 1DE. **c)** All regions except vmPFC increased mean firing rate by more than 1Hz before choice relative to fixation. Error bars represent SEM across neurons within brain region.

## Footnotes

1 pBonferroni = 0.03.

2 pBonferroni = .007

3 *p*Bonferonni = 0.012.

## References

1. Padoa-Schioppa, C. Neurobiology of Economic Choice: A Good-Based Model. Annu Rev Neurosci 34, 333–359 (2011).

2. Lee, D., Seo, H. & Jung, M. W. Neural Basis of Reinforcement Learning and Decision Making. Annu Rev Neurosci 35, 287–308 (2012).

3. Kennerley, S. W., Behrens, T. E. J. & Wallis, J. D. Double dissociation of value computations in orbitofrontal and anterior cingulate neurons. Nat Neurosci 14, 1581– 1589 (2011).

4. Klein-Flügge, M. C., Bongioanni, A. & Rushworth, M. F. S. Medial and orbital frontal cortex in decision-making and flexible behavior. Neuron vol. 110 2743–2770 Preprint at 10.1016/j.neuron.2022.05.022 (2022).

5. Montague, P. R. & Berns, G. S. Neural economics and the biological substrates of valuation. Neuron 36, 265–284 (2002).

6. Ballesta, S., Shi, W., Conen, K. E. & Padoa-Schioppa, C. Values encoded in orbitofrontal cortex are causally related to economic choices. Nature 588, 450–453 (2020).

7. Hunt, L. T. et al. Triple dissociation of Attention and decision computations across prefrontal cortex. Nat Neurosci 21, 1471–1481 (2018).

8. Rudebeck, P. H. et al. Frontal cortex subregions play distinct roles in choices between actions and stimuli. Journal of Neuroscience 28, 13775–13785 (2008).

9. Walton, M. E., Behrens, T. E. J., Buckley, M. J., Rudebeck, P. H. & Rushworth, M. F. S. Separable Learning Systems in the Macaque Brain and the Role of Orbitofrontal Cortex in Contingent Learning. Neuron 65, 927–939 (2010).

10. Knudsen, E. B. & Wallis, J. D. Taking stock of value in the orbitofrontal cortex. Nature Reviews Neuroscience vol. 23 428–438 Preprint at 10.1038/s41583-022-00589-2 (2022).

11. Yoo, S. B. M. & Hayden, B. Y. Economic Choice as an Untangling of Options into Actions. Neuron 99, 434–447 (2018).

12. Bongioanni, A. et al. Activation and disruption of a neural mechanism for novel choice in monkeys. Nature 591, 270–274 (2021).

13. Park, S. A., Miller, D. S. & Boorman, E. D. Inferences on a multidimensional social hierarchy use a grid-like code. Nat Neurosci 24, 1292–1301 (2021).

14. Bush, D., Barry, C., Manson, D. & Burgess, N. Using Grid Cells for Navigation. Neuron 87, 507–520 (2015).

15. Constantinescu, A. O., O’Reilly, X. J. & Behrens, T. E. J. Organising conceptual knowledge in humans with a gridlike code. Science (1979) 352, 1464–1468 (2016).

16. Behrens, T. E. J. et al. What Is a Cognitive Map? Organizing Knowledge for Flexible Behavior. Neuron 100, 490–509 (2018).

17. Foster, D. J. Replay Comes of Age. Annu Rev Neurosci 40, 581–602 (2017).

18. Ólafsdóttir, H. F., Bush, D. & Barry, C. The Role of Hippocampal Replay in Memory and Planning. Current Biology 28, R37–R50 (2018).

19. Dordek, Y., Soudry, D., Meir, R. & Derdikman, D. Extracting grid cell characteristics from place cell inputs using non-negative principal component analysis. Elife 5, 1–36 (2016).

20. Stachenfeld, K. L., Botvinick, M. M. & Gershman, S. J. The hippocampus as a predictive map. Nat Neurosci 20, 1643–1653 (2017).

21. Whittington, J. C. R. et al. The Tolman-Eichenbaum Machine: Unifying Space and Relational Memory through Generalization in the Hippocampal Formation. Cell 183, 1249–1263.e23 (2020).

22. Kurth-Nelson, Z. et al. Replay and compositional computation. Neuron 111, 454–469 (2023).

23. Aronov, D., Nevers, R. & Tank, D. W. Mapping of a non-spatial dimension by the hippocampal-entorhinal circuit. Nature 543, 719–722 (2017).

24. Khodagholy, D. & Gelinas, J. N. Learning-enhanced coupling between ripple oscillations in association cortices and hippocampus. Science (1979) 372, 369–372 (2017).

25. Liu, Y., Dolan, R. J., Kurth-Nelson, Z. & Behrens, T. E. J. Human Replay Spontaneously Reorganizes Experience. Cell (2019) doi:10.1016/j.cell.2019.06.012.

26. Park, S. A., Miller, D. S., Nili, H., Ranganath, C. & Boorman, E. D. Map Making: Constructing, Combining, and Inferring on Abstract Cognitive Maps. Neuron 107, 1226–1238.e8 (2020).

27. Baram, A. B., Muller, T. H., Nili, H., Garvert, M. M. & Behrens, T. E. J. Entorhinal and ventromedial prefrontal cortices abstract and generalize the structure of reinforcement learning problems. Neuron 109, 713–723.e7 (2021).

28. Aggleton, J. P., Wright, N. F., Rosene, D. L. & Saunders, R. C. Complementary patterns of direct amygdala and hippocampal projections to the macaque prefrontal cortex. Cerebral Cortex 25, 4351–4373 (2015).

29. Anderson, M. C., Bunce, J. G. & Barbas, H. Prefrontal–hippocampal pathways underlying inhibitory control over memory. Neurobiol Learn Mem 134, 145–161 (2016).

30. Kennerley, S. W., Dahmubed, A. F., Lara, A. H. & Wallis, J. D. Neurons in the Frontal Lobe Encode the Value of Multiple Decision Variables. 1162–1178 (2008).

31. Boorman, E. D., Rushworth, M. F. & Behrens, T. E. Ventromedial prefrontal and anterior cingulate cortex adopt choice and default reference frames during sequential multi-alternative choice. Journal of Neuroscience 33, 2242–2253 (2013).

32. FitzGerald, T. H. B., Seymour, B. & Dolan, R. J. The role of human orbitofrontal cortex in value comparison for incommensurable objects. Journal of Neuroscience 29, 8388–8395 (2009).

33. Jocham, G., Hunt, L. T., Near, J. & Behrens, T. E. J. A mechanism for value-guided choice based on the excitation-inhibition balance in prefrontal cortex. Nat Neurosci 15, 960–961 (2014).

34. Strait, C. E., Sleezer, B. J. & Hayden, B. Y. Signatures of value comparison in ventral striatum neurons. PLoS Biol 13, 1–22 (2015).

35. Monosov, I. E. & Hikosaka, O. Regionally distinct processing of rewards and punishments by the primate ventromedial prefrontal cortex. Journal of Neuroscience 32, 10318–10330 (2012).

36. Aquino, T. G., Cockburn, J., Mamelak, A. N., Rutishauser, U. & O’Doherty, J. P. Neurons in human pre-supplementary motor area encode key computations for value-based choice. Nat Hum Behav 7, 970–985 (2023).

37. Kunz, L. et al. Mesoscopic Neural Representations in Spatial Navigation. Trends Cogn Sci 23, 615–630 (2019).

38. Doeller, C. F., Barry, C. & Burgess, N. Evidence for grid cells in a human memory network. Nature 463, 657–661 (2010).

39. Maidenbaum, S., Miller, J., Stein, J. M. & Jacobs, J. Grid-like hexadirectional modulation of human entorhinal theta oscillations. PNAS 115, 10798–10803 (2018).

40. Jacobs, J. et al. Direct recordings of grid-like neuronal activity in human spatial navigation. Nat Neurosci 16, 1188–1190 (2013).

41. Joo, H. R. & Frank, L. M. The hippocampal sharp wave–ripple in memory retrieval for immediate use and consolidation. Nat Rev Neurosci 19, 744–757 (2018).

42. Fyhn, M., Hafting, T., Treves, A., Moser, M. B. & Moser, E. I. Hippocampal remapping and grid realignment in entorhinal cortex. Nature 446, 190–194 (2007).

43. Buzsàki, G. & Eidelberg, E. Phase relations of hippocampal projection cells and interneurons to theta activity in the anesthetized rat. Brain Res 266, 334–339 (1983).

44. Knudsen, E. B. & Wallis, J. D. Closed-Loop Theta Stimulation in the Orbitofrontal Cortex Prevents Reward-Based Learning. Neuron 1–11 (2020) doi:10.1016/j.neuron.2020.02.003.

45. Strait, C. E., Blanchard, T. C. & Hayden, B. Y. Reward value comparison via mutual inhibition in ventromedial prefrontal cortex. Neuron 82, 1357–1366 (2014).

46. Johnson, A. & Redish, A. D. Neural Ensembles in CA3 Transiently Encode Paths Forward of the Animal at a Decision Point. Journal of Neuroscience 27, 12176–12189 (2007).

47. Vaz, A. P., Inati, S. K. & Zaghloul, K. A. Replay of cortical spiking sequences during human memory retrieval. Science (1979) 1134, 1131–1134 (2020).

48. Vaz, A. P., Inati, S. K., Brunel, N. & Zaghloul, K. A. Coupled ripple oscillations between the medial temporal lobe and neocortex retrieve human memory. Science (1979) 363, 975–978 (2019).

49. Norman, Y. et al. Hippocampal sharp-wave ripples linked to visual episodic recollection in humans. Science (1979) 365, (2019).

50. Karlsson, M. P. & Frank, L. M. Awake replay of remote experiences in the hippocampus. Nat Neurosci 12, 913–918 (2009).

51. Logothetis, N. K. et al. Hippocampal-cortical interaction during periods of subcortical silence. Nature 491, 547–553 (2012).

52. Liu, A. A. et al. A consensus statement on detection of hippocampal sharp wave ripples and differentiation from other fast oscillations. Nature Communications vol. 13 Preprint at 10.1038/s41467-022-33536-x (2022).

53. Ramirez-villegas, J. F., Logothetis, N. K. & Besserve, M. Diversity of sharp-wave – ripple LFP signatures reveals differentiated brain-wide dynamical events. PNAS (2015) doi:10.1073/pnas.1518257112.

54. Michon, F., Sung, J.-J., Kim, Y. C., Ciliberti, D. & Kloosterman, F. Post-learning Hippocampal Replay Selectively Reinforces Spatial Memory for Highly Rewarded Article Post-learning Hippocampal Replay Selectively Reinforces Spatial Memory. Current Biology 29, 1436–1444 (2019).

55. Buzsaki, G. Hippocampal Sharp Wave-Ripple : A Cognitive Biomarker for Episodic Memory and Planning. Hippocampus 1188, 1073–1188 (2015).

56. Yu, J. Y. & Frank, L. M. Hippocampal-cortical interaction in decision making. Neurobiol Learn Mem 117, 34–41 (2015).

57. Rangel, A., Camerer, C. & Montague, P. R. A framework for studying the neurobiology of value-based decision making. Nat Rev Neurosci 9, 545–556 (2008).

58. Padoa-Schioppa, C. & Cai, X. The orbitofrontal cortex and the computation of subjective value: Consolidated concepts and new perspectives. Ann N Y Acad Sci 1239, 130–137 (2011).

## Supplementary References

1. Hunt, L. T. et al. Triple dissociation of Attention and decision computations across prefrontal cortex. Nat Neurosci 21, 1471–1481 (2018).

2. Butler, J. L., et al. Covert valuation for information sampling and choice. Biorxiv (2021) doi:10.1101/2021.10.08.463476.

3. Cavanagh, S. E., Malalasekera, W. M. N., Miranda, B., Hunt, L. T. & Kennerley, S. W. Visual fixation patterns during economic choice reflect covert valuation processes that emerge with learning. Proc Natl Acad Sci U S A 116, 22795–22801 (2019).

4. Muller, T. H., et al. Distributional reinforcement learning in prefrontal cortex. Biorxiv (2021) doi:10.1101/2021.06.14.448422.

5. Oostenveld, R., Fries, P., Maris, E. & Schoffelen, J. FieldTrip : Open Source Software for Advanced Analysis of MEG, EEG, and Invasive Electrophysiological Data. 2011, (2011).

6. Doeller, C. F., Barry, C. & Burgess, N. Evidence for grid cells in a human memory network. Nature 463, 657–661 (2010).

7. Constantinescu, A. O., O’Reilly, X. J. & Behrens, T. E. J. Organising conceptual knowledge in humans with a gridlike code. Science (1979) 352, 1464–1468 (2016).

8. Chen, D. et al. Theta oscillations coordinate grid-like representations between ventromedial prefrontal and entorhinal cortex. Sci Adv 7, 1–13 (2021).

9. Maidenbaum, S., Miller, J., Stein, J. M. & Jacobs, J. Grid-like hexadirectional modulation of human entorhinal theta oscillations. PNAS 115, 10798–10803 (2018).

10. Prelec, D. The Probability Weighting Function. vol. 66 (1998).

11. Bongioanni, A. et al. Activation and disruption of a neural mechanism for novel choice in monkeys. Nature 591, 270–274 (2021).

12. Imaizumi, Y., Tymula, A., Tsubo, Y., Matsumoto, M. & Yamada, H. A neuronal prospect theory model in the brain reward circuitry. Nat Commun 13, (2022).

13. Schwarz, G. Estimating the Dimension of a Model. Ann Stat 6, 461–464 (1978).

14. Park, S. A., Miller, D. S. & Boorman, E. D. Inferences on a multidimensional social hierarchy use a grid-like code. Nat Neurosci 24, 1292–1301 (2021).

15. Fyhn, M., Hafting, T., Treves, A., Moser, M. B. & Moser, E. I. Hippocampal remapping and grid realignment in entorhinal cortex. Nature 446, 190–194 (2007).

16. Logothetis, N. K. et al. Hippocampal-cortical interaction during periods of subcortical silence. Nature 491, 547–553 (2012).

17. Vaz, A. P., Inati, S. K., Brunel, N. & Zaghloul, K. A. Coupled ripple oscillations between the medial temporal lobe and neocortex retrieve human memory. Science (1979) 363, 975–978 (2019).

18. Karlsson, M. P. & Frank, L. M. Awake replay of remote experiences in the hippocampus. Nat Neurosci 12, 913–918 (2009).

19. Liu, A. A. et al. A consensus statement on detection of hippocampal sharp wave ripples and differentiation from other fast oscillations. Nature Communications vol. 13 Preprint at 10.1038/s41467-022-33536-x (2022).

20. Staudigl, T. et al. Hexadirectional Modulation of High-Frequency Electrophysiological Activity in the Human Anterior Medial Temporal Lobe Maps Visual Space. Current Biology 28, 3325–3329.e4 (2018).

21. Kahneman, D. & Tversky, A. Prospect Theory: An Analysis of Decision under Risk. Econometrica 47, 263–291 (1979).

22. Lakshminarayanan, V. R., Chen, M. K. & Santos, L. R. The evolution of decision-making under risk: Framing effects in monkey risk preferences. J Exp Soc Psychol 47, 689–693 (2011).

23. Jacobs, J. et al. Direct recordings of grid-like neuronal activity in human spatial navigation. Nat Neurosci 16, 1188–1190 (2013).

